# Replication Fork-Associated Checkpoint Mediator Mrc1/Claspin Acts in Arrest-Independent Double-Strand Break Survival

**DOI:** 10.64898/2026.09.15.751834

**Authors:** Marissa M. Ashton, Elliot Busch, James E. Haber

## Abstract

In budding yeast, the DNA damage checkpoint kinase Mec1^ATR^ is the primary sensor of DSB-associated single-stranded DNA and coordinates cell cycle arrest with DNA repair. Deletion of Mec1 completely abolishes G2/M cell cycle arrest; however, approximately half of *mec1*Δ cells still survive an endonuclease-induced DSB, repairing the DSB either by single-strand annealing or break-induced replication. Here we show that Mec1-independent DSB repair is independent of Rad9 but requires the Tel1^ATM^ kinase, Rad53 kinase, and the 9-1-1 sliding clamp, which promotes Tel1 retention at damage sites. We identify Mrc1^Claspin^, a replication fork-associated checkpoint mediator, as an unexpected contributor to DSB survival in the absence of Mec1. Unlike Rad9, the canonical DNA damage adaptor, Mrc1’s role in DSB survival is independent of its Mec1/Tel1 consensus phosphorylation sites and relies primarily on its C-terminal domain. Mrc1 accumulates at a single DSB and both promotes end-tethering and limits DNA end-resection in an S-phase-specific manner. Mrc1 functions independently of its replication checkpoint partners, Tof1 and Csm3. We further show that heterochromatic gene silencing in budding yeast is Mrc1-dependent but Tof1– and Csm3-independent. Together, these findings define a Mec1-independent survival pathway and establish Mrc1 as a novel regulator of DSB repair.

## Introduction

DNA double-strand breaks (DSBs) threaten genomic stability and cell viability. These lesions can arise from exogenous sources such as ionizing radiation or from endogenous processes including replication fork stalling and replication stress. Cells have evolved sensitive checkpoint mechanisms to detect DSBs and trigger cell cycle arrest to provide time for repair. Understanding these pathways is essential for understanding how genomic integrity is maintained and how defects in checkpoint regulation contribute to genomic instability. In the budding yeast *Saccharomyces cerevisiae*, the primary DSB checkpoint is mediated by the Mec1 ^ATR^ kinase and to a much lesser extent by its paralog, Tel1^ATM^. Mec1-Ddc2^ATRIP^ is recruited by RPA-coated single-stranded DNA (ssDNA) created by 5’ to 3’ resection at DSBs. In addition, the 9-1-1 clamp (Rad17, Mec3, and Ddc1), which is loaded at single– stranded/double-stranded DNA junctions, amplifies the Mec1 checkpoint signal. Mec1 phosphorylates numerous substrates including the adaptor protein Rad9 and the effector kinase Rad53 (reviewed in 1,2). This signaling cascade leads to G2/M cell cycle arrest, stabilization of repair intermediates, and coordination with DNA repair machinery (3,4,5). Tel1 kinase, while phosphorylating many of the same substrates as Mec1, plays a limited role in the canonical DSB response. Tel1 is recruited to unresected DSB ends by the MRX complex (Mre11-Rad50-Xrs2), which directly binds DNA ends. The Mre11-Rad50 subcomplex, when in its ATP-bound closed conformation, physically bridges the two ends of the DSB—a process termed end-tethering—and holds broken ends in close proximity (6,7). End tethering is critical for timely and faithful DSB repair, as it maintains the spatial proximity necessary for accurate repair and limits the risk of aberrant genomic rearrangements (8).

Previous work has shown that when a DSB can be repaired, by either single-strand annealing (SSA) or break-induced replication (BIR), approximately 50% of *mec1*Δ cells can survive, as measured by colony forming units, where at least one offspring of a damaged cell can complete repair (9,10). This observation raises fundamental questions: What molecular machinery enables Mec1-independent survival? How is repair accomplished in the absence of checkpoint-induced arrest? What role do other checkpoint components play when Mec1 is absent?

In addition to its role in DSB checkpoints, Mec1 is the primary mediator of the inter-S phase checkpoint, which is activated by stalled replication forks (reviewed in 11). Mrc1^Claspin^ is an evolutionarily conserved replication fork protein that mediates replication fork checkpoint signaling, coupling increased ssDNA at stalled forks to Mec1 activation (12,13). Mec1 activates Rad53, which inhibits firing of late replication origins (Diffley,Shirahige). Mrc1 associates with Tof1 and Csm3 at replication forks and is phosphorylated by Mec1 at multiple sites (14). Unlike *tof1*Δ cells, *mrc1*Δ cells exhibit measurably slowed DNA replication, indicating that Mrc1 plays a more direct role in replication fork progression (15).

Mrc1 extends across one face of the CMG (Cdc45-Mcm2-7-GINS) helicase, with its N-terminus contacting Tof1-Csm3, its central region linked to Mcm2 and Mcm6, and its C-terminal residues in proximity to Cdc45, positioning Mrc1 to coordinate helicase and leading-strand polymerase activities across the replisome (16). Additionally, the central domain of Mrc1 (residues 433–457) mediates a direct physical interaction with Cdc45 that is required to maintain Cdc45 and Pol2 localization at stalled replication forks (17). Mrc1 and Cdc45 are thought to function in parallel, both acting as recruiters of the effector kinase Rad53 to the stalled replisome: Mrc1 is modified by Mec1 and Rad53-dependent phosphorylation, while Cdc45 is mostly phosphorylated by Rad53 (16).

Mrc1 is predicted to interact with Ctf4, a homotrimeric replisome scaffold that bridges the CMG helicase and DNA polymerase α. Ctf4 also recruits the Chl1 helicase to the fork via a conserved CIP box motif (18,19). Chl1^ChlR1/Ddx11^ in turn physically contacts the cohesin ring to facilitate its acetylation and stabilization during S phase, thereby coupling DNA replication with the establishment of sister chromatid cohesion (18). Mrc1 was shown to play a role in limiting resection at stalled replication forks, in the presence of hydroxy urea (HU), but also at an endonuclease-induced DSB (20,21). Mrc1 also appears to promote chromatin compaction after HU addition (21). Whether Mrc1 has further roles in DSB repair has not been fully explored.

Here, we investigated the Mec1-independent DSB survival pathway and identified unexpected roles for both Mrc1 and Ctf4 in DSB repair. Our findings establish a novel, Rad9-independent checkpoint pathway requiring Tel1, Rad53, and the 9-1-1 complex, and reveal that Mrc1 contributes to DSB survival through its functions in broken end-tethering and in limiting 5’ to 3’ resection during S phase.

## Results

### Mec1 is required for DSB-induced arrest but not for DSB survival

To investigate DSB checkpoint signaling and repair, we used a system in which a galactose–induced HO endonuclease creates a single DSB on the left arm of chromosome III, between flanking homologous 1-kb repeats, one of which is located 20 kb from the DSB (10,8). This DSB can be repaired either through single-strand annealing (SSA) or break-induced replication (BIR) (Fig. 1A and Supplemental Figure 1) (reviewed in 22). Both SSA and BIR require at least 5-6 hr to repair the break. In the case of SSA, this time is required for 5’ to 3’ resection, which occurs at approximately 4 kb/hr, to expose the distant flanking homology (10). In the case of BIR, this delay is attributable to a Recombination Execution Checkpoint (REC), a surveillance mechanism that monitors whether the two DSB ends can synapse with homologous sequences in a productive orientation before facilitating the initiation of repair DNA synthesis (23,8). When only one end is available for strand invasion (as in BIR) the REC imposes an Sgs1– and Mph1-dependent delay in repair commitment. After DSB induction, cells accumulate at G2/M with a characteristic dumbbell-shape and a single nucleus, visible by DAPI staining. Arrest persists for about 6 h, when repair is accomplished (Fig. 1B). In contrast, *mec1*Δ cells fail to arrest and continue through the cell cycle with a doubling time equivalent to cells without an induced DSB, confirming that Mec1 is the sole checkpoint kinase required for DSB-induced arrest (Fig. 1B). Despite failing to arrest, and the fact that repair takes almost three cell division times to be completed, *mec1*Δ cells survive DSB induction with approximately 50% wild-type viability, as measured by colony-forming units (Fig. 1C), revealing a checkpoint arrest-independent survival mechanism (9). When repair is measured by PCR, it becomes evident that only about 25% of broken chromosomes are repaired (Fig. 1D). Repair can be substantially improved by artificially arresting cells in G2/M by the addition of nocodazole (Fig. 1D).

**Figure 1.**
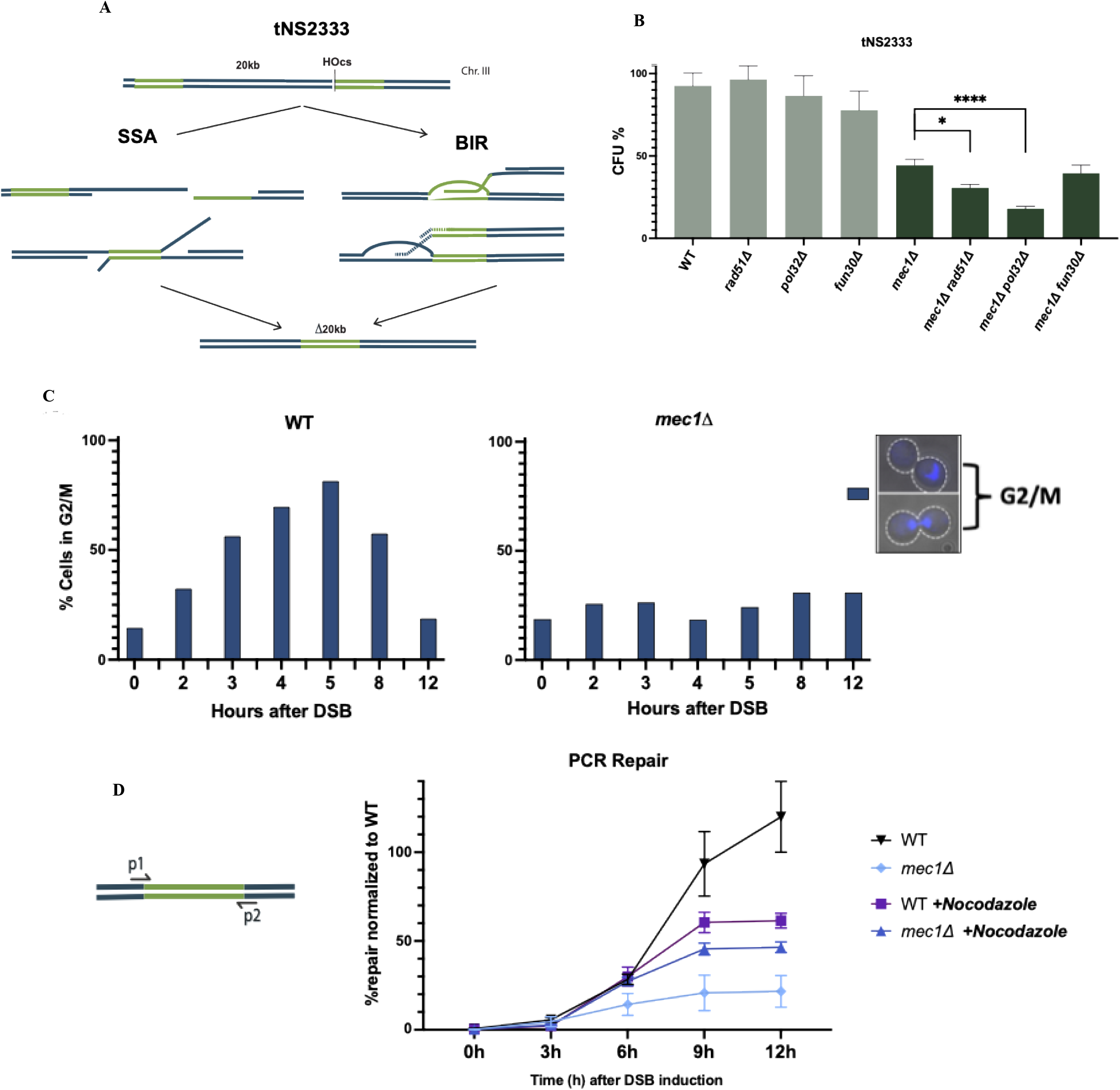
Mec1 is required for checkpoint arrest but not for DSB survival. A. Schematic of repair in strain tNS2333. A galactose-inducible HO endonuclease generates a site-specific DSB at LEU2 on chromosome III, flanked by a homologous sequence 20 kb away. The break can be repaired by single-strand annealing (SSA) or break-induced replication (BIR). B. Cell-cycle morphology (DAPI staining) of wild-type and *mec1*Δ cells at the indicated times after DSB induction. Wild-type cells arrest in G2/M for up to 8 h before completing repair; *mec1*Δ cells show no detectable arrest. C. Colony-forming unit (CFU) assay measuring DSB survival (galactose-induced/uninduced ratio) in wild-type, *mec1*Δ*, rad51*Δ, *pol32*Δ, *fun30*Δ single, double and triple mutants. Approximately 50% of *mec1*Δ cells survive despite the absence of checkpoint arrest. *mec1*Δ *rad51*Δ and *mec1*Δ *pol32*Δ drop to 25% CFU or less, while *mec1*Δ *fun30*Δ looks like *mec1*Δ. D. PCR-based repair assay in strain tNS2333 showing primers designed around the repair product only. Quantification of repair kinetics. X-axis is time after DSB induction. The y-axis is normalized to the PCR signal % of a 100% repaired sample (WT colony grown on Yp-gal) relative to 0 h PCR signal for the repair product normalized to a control amplicon at *GLC7*. *mec1*Δ cells show defect in repair, which is rescued by arresting cells with Nocodazole.

While we had previously assumed that repair in this system proceeded principally through SSA (10,8), we now understand that Mec1-independent survival can occur also through BIR. To distinguish between these two processes, we examined mutations affecting one or the other process. SSA is a Rad51-independent process but highly dependent on resection, which is carried out by many factors including the chromatin remodeler Fun30 (24,Chen X et al., 2012,25). A *mec1*Δ *fun30*Δ strain shows no additional defect, implying that disrupting the SSA pathway is not crucial for Mec1-independent survival. In contrast, *mec1*Δ *rad51*Δ cells show reduced viability, as measured by colony forming units (CFU), compared to *mec1*Δ single mutants. Similarly, BIR depends on Pol32 (26) and *mec1*Δ *pol32*Δ cells also show a marked reduction in repair. Wild-type cells can survive with either repair pathway individually disrupted (Fig. 1C). Interestingly, there are still cells in a *mec1*Δ *pol32*Δ *fun30*Δ population that are able to survive the DSB.

### Mec1-independent survival relies on Rad53 and Tel1, which is retained at the DSB by the 9-1-1 complex

Mec1-independent repair still requires checkpoint signaling components. Both Tel1 and Rad53 are essential: *mec1*Δ *tel1*Δ double mutants exhibit significantly reduced viability after DSB induction, while *mec1*Δ *rad53*Δ cells are almost completely inviable (Fig. 2A). *mec1*Δ *mre11*Δ cells repair poorly. This reduction could reflect the role of the MRX complex in recruiting Tel1 to the DSB (4), but may also implicate Mre11’s roles in promoting end resection or end-tethering (27,28).

**Figure 2.**
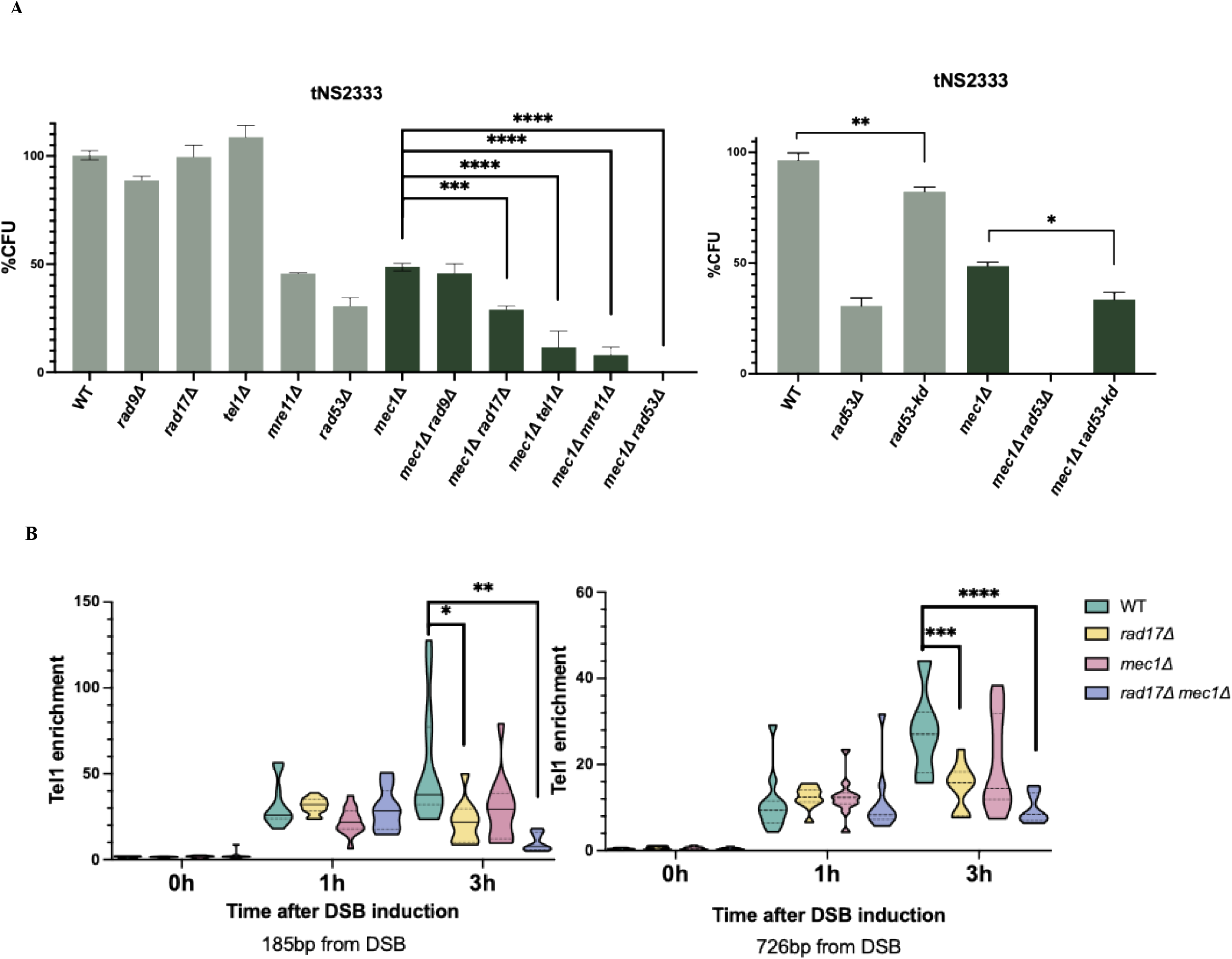
Mec1-independent DSB survival requires Tel1, Rad53, and the 9-1-1 complex. A. CFU assay in strain tNS2333. Deletion of Tel1, Rad53, or Rad17 in combination with *mec1*Δ abrogates Mec1-independent survival, whereas deletion of Rad9 has no effect. Rad53 kinase activity is dispensable for DSB survival. B. ChIP of HA-Tel1 at 726 bp and 185 bp from the DSB, x-axis is time after DSB induction, y-axis is fold ChIP enrichment relative to an uncut locus and normalized to input. Median shown in dashed horizontal line, quartile shown in dotted horizontal line.

To understand how Rad53 acts in Mec1-independent DSB survival, we examined its role as a protein kinase (Fig. 2B). Notably, a kinase-dead Rad53 (*rad53-K227A*) (29,30) only mildly impacts survival in general, indicating that Rad53 could function as an adaptor or structural protein rather than through its kinase activity in this context. However, the kinase-dead mutation resembles both *mec1*D and *rad53*D when we measure its effect on G2/M arrest (Fig. S3). In *mec1*Δ *rad17*Δ cells, where the 9-1-1 complex is disrupted, there is a more moderate, but still significant, reduction in viability (Fig. 2A). Loss of kinases Chk1 or Dun1 in *mec1*Δ cells showed no impact on Mec1-independent survival (Fig. S2).

Chromatin immunoprecipitation (ChIP) revealed that Tel1 accumulates near DSB sites in both wild-type and *mec1*Δ cells; critically, this accumulation is diminished in *rad17*Δ cells but not significantly in *mec1*Δ strains (Fig. 2C). This result suggests that the 9-1-1 complex retains Tel1 at damage sites, providing a mechanism for 9-1-1 function in Mec1-independent DSB repair. In an attempt to further activate the Tel1-specific checkpoint in *mec1*Δ cells, we tested the hypermorphic tel1-*hy909* allele that has been shown to compensate for the lack of Mec1 function (when/where?) through increased Tel1 kinase activity and association with DSBs (31). However, tel1-*hy909* showed no significant improvement in Mec1-independent survival (Fig. S4). Since one of Tel1’s substrates is histone H2A, we also tested whether Mec1-independent survival requires histone H2A phosphorylation by Tel1. Non-phosphorylatable or phosphomimetic histone H2A alleles at serine 129 (*H2A-S129A* or *H2A-S129E*) had no effect on Mec1-independent survival (Fig. S4). Taken together, these results imply that although Tel1 is crucial for Mec1-independent survival, hyperactive Tel1 cannot rescue survival in this system, and Tel1’s role does not rely on phosphorylation of H2A-S129.

### Mrc1 promotes DSB survival independently of its replication checkpoint function

Having established that the Tel1/Rad53/9-1-1 pathway promotes survival in the absence of Mec1, we sought additional factors contributing to Mec1-independent survival. Unlike Rad9, which showed little importance in the Mec1-independent pathway, Mrc1, known as Rad9’s counterpart in the replication fork checkpoint32, emerged as a critical factor (Fig. 3). Whereas a *mrc1*Δ single mutant showed a modest but significant defect in DSB survival, *mec1*Δ *mrc1*Δ double mutants showed more significantly reduced viability compared to *mec1*Δ (Fig. 3A). Mrc1’s partners Csm3 and Tof1, showed more modest defects in the Mec1-independent pathway (Fig. 3a). We found a similar defect when Ctf4 was deleted, both in Mec1^+^ cells and in in the absence of Mec1.

**Figure 3.**
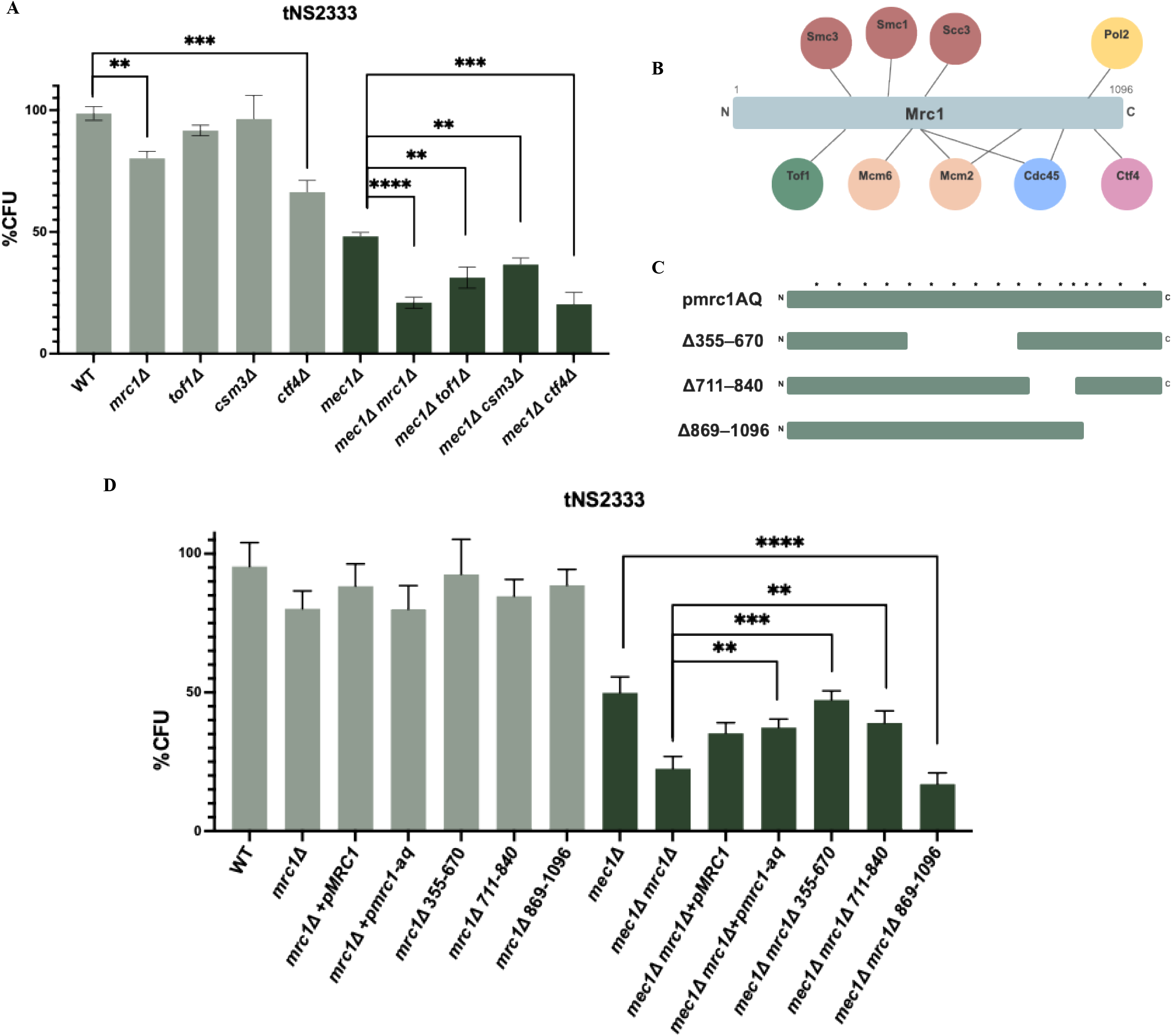
Mrc1 is required for Mec1-independent DSB survival through its C-terminal domain. A. CFU assay in strain tNS2333. Mrc1, Tof1, Csm3, and Ctf4 each contribute to Mec1-independent DSB survival. B. Schematic of Mrc1 and its predicted physical interactors (adapted from Shrestha et al., 2023). C. Diagram of Mrc1 domain deletion and phosphorylation-site mutant constructs. D. CFU assay in *mec1*Δ *mrc1*Δ cells complemented with wild-type *MRC1* or *mrc1-AQ* (17 SQ/TQ→AQ; Mec1 phosphorylation sites ablated). *mrc1-AQ* rescues survival equivalently to wild-type. *mrc1*Δ*355–670* and *mrc1*Δ*711–840* do not impair Mec1-independent DSB survival. *mrc1*Δ*869–1096* phenocopies *mrc1*Δ, mapping the essential function to the Mrc1 C-terminal domain.

In the intra-S checkpoint, phosphorylation of Mrc1 at 17 S/T Q sites by Mec1 is a key aspect of regulation (13). Remarkably, a plasmid carrying *mrc1-aq*, lacking these phosphorylation sites, rescued *mec1*Δ *mrc1*Δ lethality as effectively as a wild-type copy of Mrc1. We examined several previously characterized deletions of Mrc1 (33,34). Deletion of the C-terminal domain (*mrc1*Δ869–1096) phenocopied *mrc1*Δ. (Fig. 3D) Deletion of Mrc1’s histone binding domain (*mrc1*Δ711–840) or deletion of much of its predicted cohesin and Cdc45 binding domains (*mrc1*Δ355–670) did not impair survival (Fig. 3D). A *mrc1-8D* phosphomimetic allele, which mimics phosphorylation of Mrc1 by Rad53 and leads to slower DNA replication (33), created a partial defect, potentially supporting a role for Mrc1 that is independent of Mec1-dependent DNA damage checkpoint signaling (Fig. S5). These results demonstrate that Mrc1 promotes DSB survival through its C-terminal domain, apparently independent of its canonical replication checkpoint function, but possibly through its interaction with Ctf4.

### Mrc1 promotes DSB repair

We then examined how repair kinetics in strain tNS2333 were affected by Mrc1, notably in *mec1*Δ *mrc1*Δ and *mec1*Δ *tel1*Δ double mutants. As noted above, using a PCR-based assay, we observed that *mec1*Δ cells repair at a much lower frequency than wild type (Fig. 1D), seen previously by Southern blot (9). While *mrc1*Δ and *tel1*Δ single mutants show no significant difference in repair kinetics from wild type, *mec1*Δ *mrc1*Δ and *mec1*Δ *tel1*Δ double mutants show significantly less repair compared to the *mec1*Δ single mutant (Fig. 4A). These repair kinetics demonstrate that Tel1 and Mrc1 promote the progression of DSB repair in the absence of canonical Mec1-dependent checkpoint activation and arrest. As previously reported for a similar SSA/BIR system (YMV80), repair can be improved by arresting cells with nocodazole (9). We recapitulated this rescue in G2/M-arrested cells in several single and double mutants (Fig. S6). We note that overall repair in nocodazole-arrested cells is notably lower than in cycling cells.

**Figure 4.**
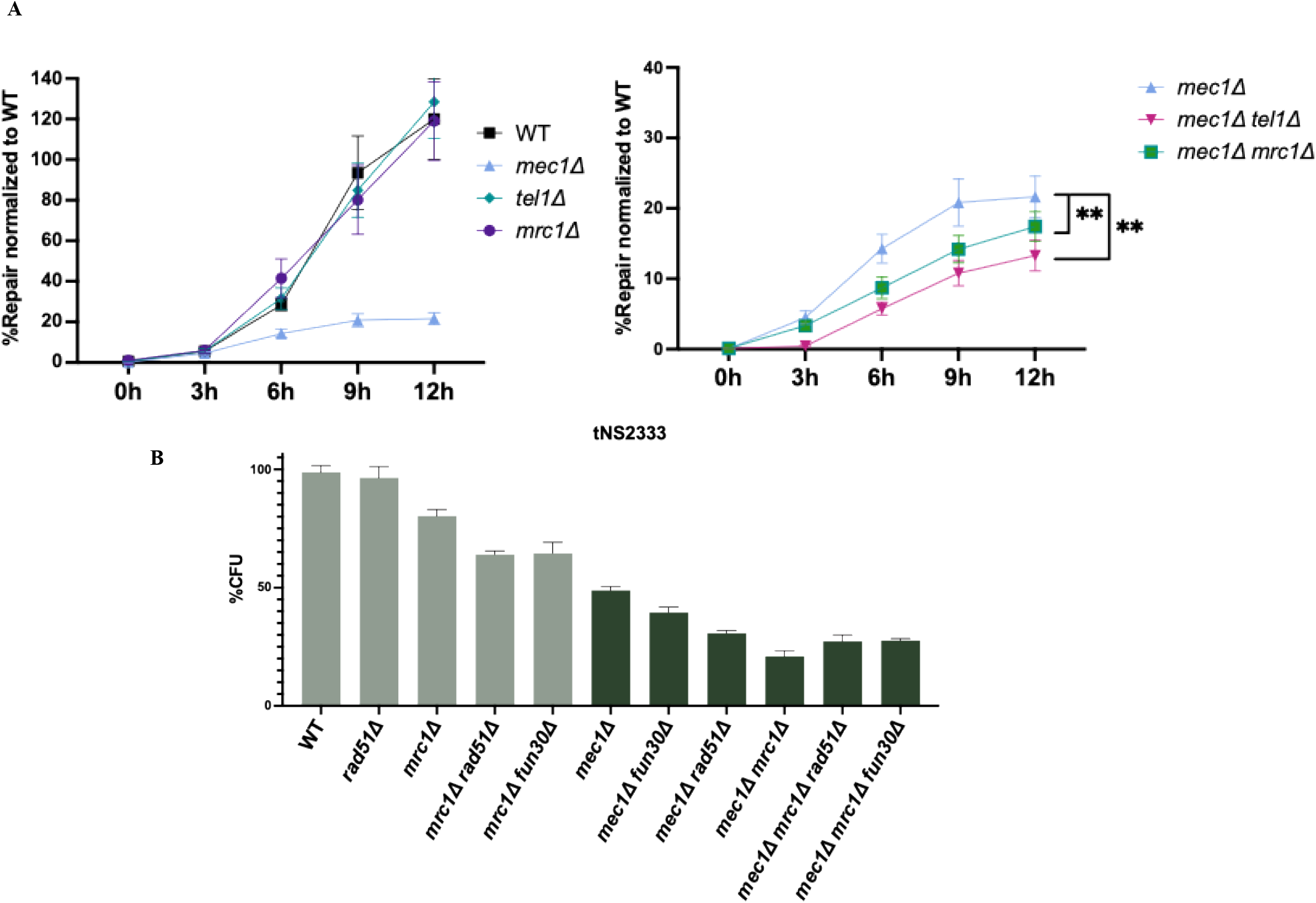
Mrc1 and Tel1 promote Mec1-independent break repair. A. Quantification of repair kinetics. X-axis is time after DSB induction. The y-axis is normalized to the PCR signal % of a 100% repaired sample (WT colony grown on Yp-gal) relative to 0 h PCR signal for the repair product normalized to a control amplicon at *GLC7*. *mec1*Δ cells show defect in repair, *mec1*Δ *tel1*Δ and *mec1*Δ *mrc1*Δ mutants show further repair defects whereas *tel1*Δ and *mrc1*Δ show no significant impact on repair. B. CFU assay in strain tNS2333. *mec1*Δ *mrc1*Δ DSB survival is not further reduced in *mec1*Δ *mrc1*Δ *rad51*Δ and *mec1*Δ *mrc1*Δ *fun30*Δ, indicating *MRC1* is epistasic to *RAD51* in the *mec1*Δ background.

Although we had shown that *mec1*D *rad51*D was more severely affected than *mec1*D *fun30*D (Fig. 2), there was no such distinction in *mec1*Δ *mrc1*Δ *rad51*Δ and *mec1*Δ *mrc1*Δ *fun30*Δ triple mutants (Fig. 4B). We then asked if Mrc1 was important in other DSB repair systems. In strain JLR092, where only BIR can repair the DSB (Fig. 5A), deleting Mrc1 markedly reduced repair in an otherwise wild-type strain and nearly eliminated repair when Mec1 was also deleted. Notably, neither Tof1 nor Csm3 is required for BIR-mediated repair (Fig. 5B). In YJK17 cells, where only interchromosomal gene conversion (GC) is available for HO-induced DSB repair (Fig. 5C, *mrc1*Δ showed a significant reduction in DSB survival, and *mec1*Δ *mrc1*Δ cells exhibited an equivalently reduced GC efficiency compared to *mec1*D (Fig. 5D). Finally, we examined mating-type switching where an HO-induced DSB at *MAT***a** is repaired using the donor *HML*a-inc (Fig. 5E). In this system, where pairing between *HML* and *MAT* is facilitated by a recombination enhancer (35), DSB repair is rapid and efficient, and the DNA checkpoint is not activated (36). Here, *mec1*Δ*, mrc1*Δ, and *tel1*Δ single mutants have no impact on survival, but, surprisingly, the double mutants *mec1*Δ *mrc1*Δ and *mec1*Δ *tel1*Δ show significant defects (Fig. 5F).

**Figure 5.**
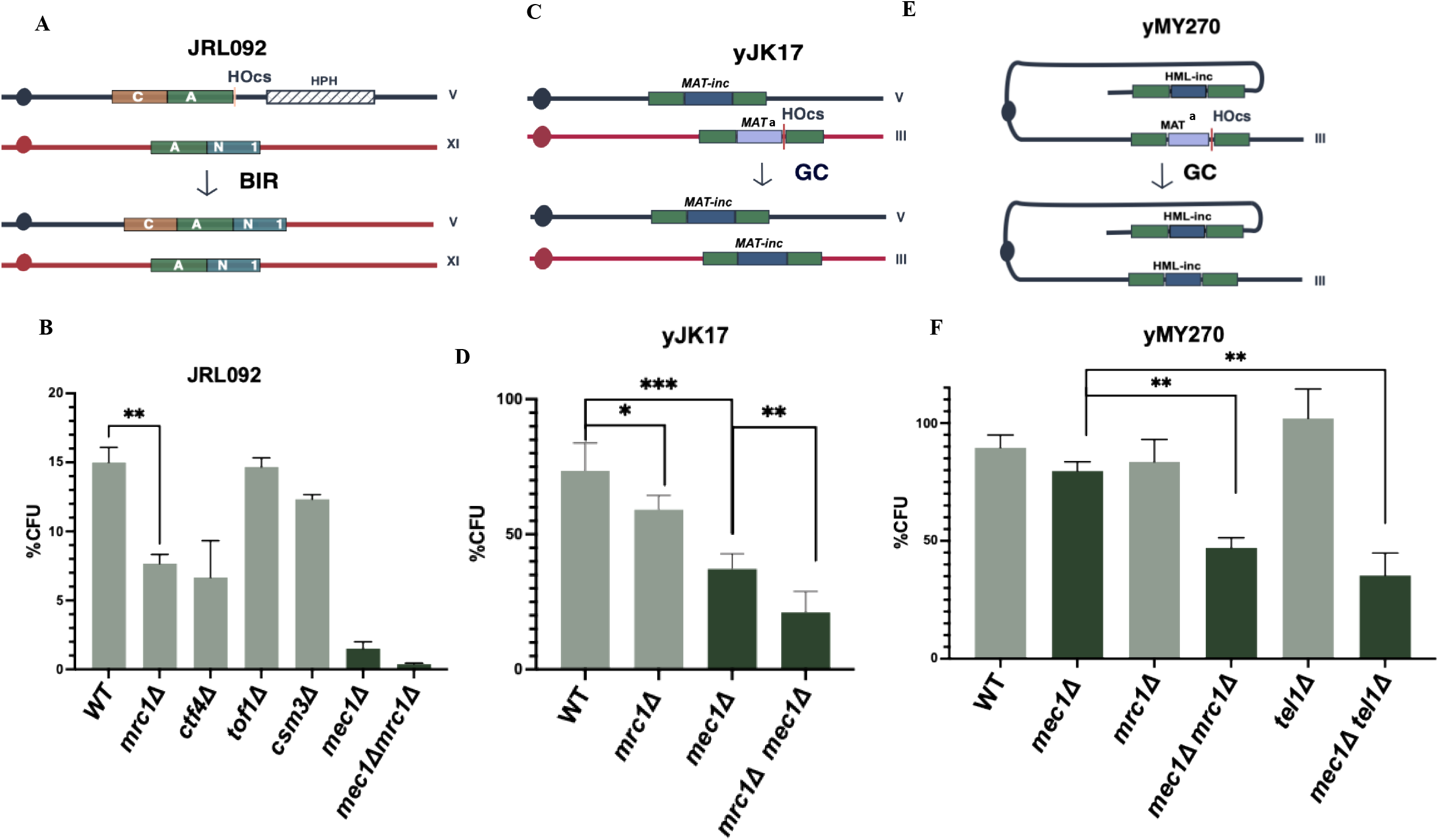
Mrc1 promotes DSB survival across multiple repair contexts. A. BIR repair schematic in strain JRL092. A galactose-inducible one-ended DSB on chromosome V is repaired by BIR using homology on chromosome XI. B. CFU assay measuring DSB survival in the indicated genotypes in strain JRL092. *mrc1*Δ but not *tof1*Δ nor *csm3*Δ show significant defects while *mec1*Δ shows a severe defect. C. Gene conversion repair schematic in strain yJK17. A galactose-inducible DSB at MAT on chromosome III is repaired using a homologous MAT-inc donor on chromosome V. D. CFU assay measuring DSB survival in the indicated genotypes in strain yJK17, *mec1*Δ *mrc1*Δ shows significant defects compared to *mec1*Δ. E. Gene conversion repair schematic for strain yMY270. A galactose-inducible DSB at MAT on chromosome III is repaired using an endogenous HML-inc donor, also on chromosome III. F. CFU assay measuring DSB survival in the indicated genotypes in strain yMY270, *mec1*Δ *mrc1*Δ and *mec1*Δ *tel1*Δ show significant defects compared to *mec1*Δ which shows no significant defects.

### Mrc1 promotes end tethering

Mrc1 and Ctf4 both interact, directly or indirectly, with cohesin, which has been implicated in DSB end tethering (28) and in facilitating Tel1-dependent phosphorylation of histones surrounding a DSB (37). In a donorless strain in which an HO-induced DSB at *MAT* is not repaired by homologous recombination, the presence of fluorescently tagged TetO:TetR-GFP and LacO:LacI-mCherry at sites flanking the DSB (28) allow quantification of DSB end-tethering (Fig. 6A). In wild-type strains nearly all ends remain tethered, and this tethering is not dependent on Mec1. In contrast, deletion of Mrc1 caused untethering in more than 20% of cells. Deletion of Tof1 had a much less profound effect (Fig. 6B). Deletions of different Mrc1 domains revealed that removing the predicted cohesin and Cdc45 binding domain (*mrc1*Δ355–670) or the C-terminal region that interacts with Ctf4 (*mrc1*Δ869–1096) each had a significant effect on end-tethering, though neither was as severe as the full deletion (Fig. 6B). These results imply that multiple domains within Mrc1 contribute to end tethering.

**Figure 6.**
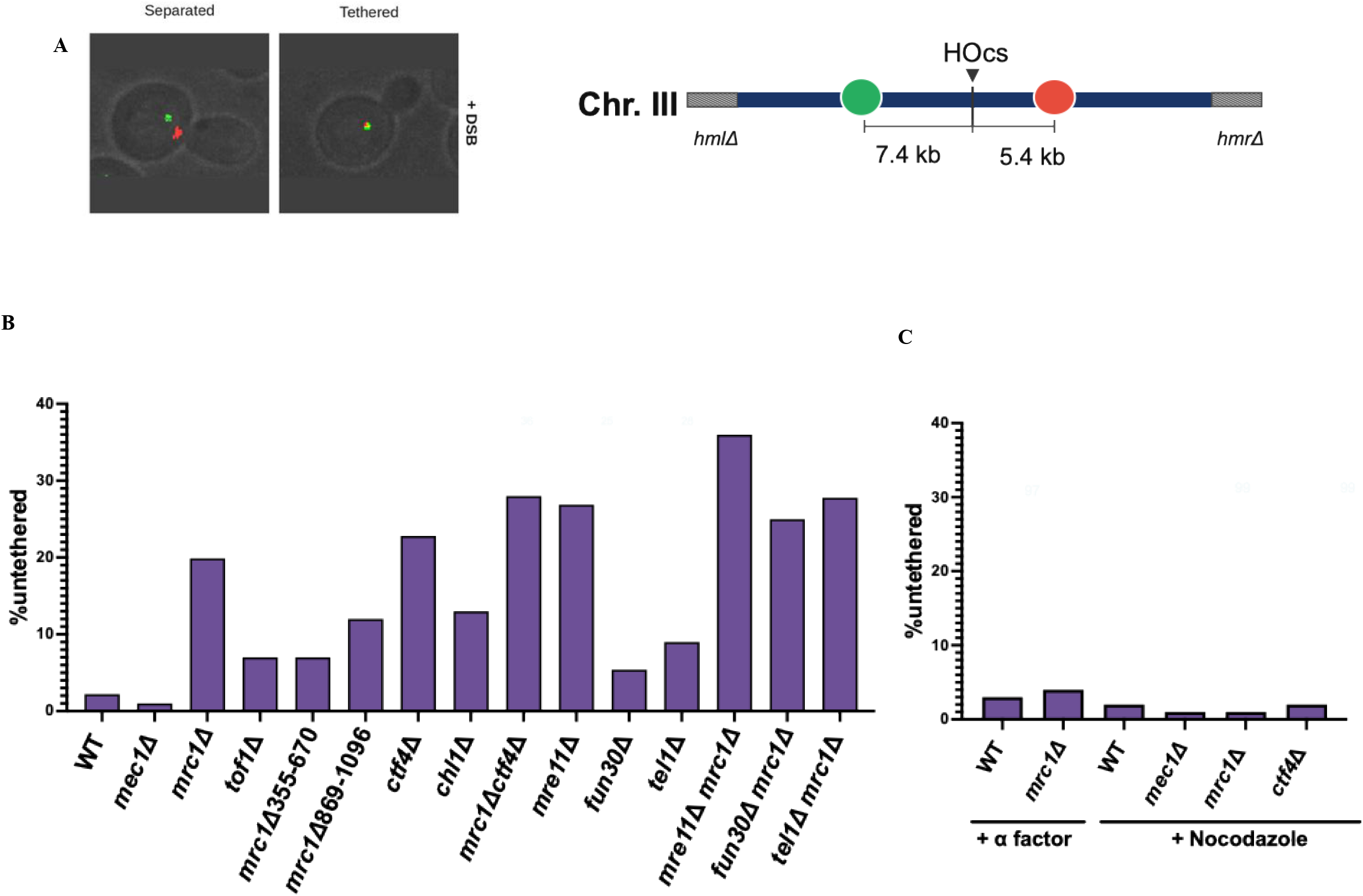
**Mrc1 promotes end tethering independently of Tof1, during S-phase**. A. End-tethering assay schematic (adapted from Phipps et al., 2025). A DSB is induced at *MAT* on chromosome III, flanked by tetO and lacO arrays bound by TetR-GFP and LacI-mCherry, respectively. Co-localization indicates intact tethering; separated foci indicate tethering failure. Representative images are shown. B. Quantification of end tethering in the indicated strains 2.5 h after DSB induction. C. *mrc1*Δ end tethering defect is abolished by G1 arrest (alpha factor) or G2/M arrest (nocodazole).

Both *ctf4*D and *chl1*D increase the loss of end-tethering (Fig. 6). The *mrc1*D *ctf4*D double mutant resembled each single mutant. As previously noted, deleting Mre11 shows a large increase in untethering (38,9,28). The effect of deleting both Mre11 and Mrc1 resulted in a significantly higher level of untethering. Deleting Tel1 had a significant increase compared to wild type, but much less than Mrc1.

Moreover, we found that the defects in end-tethering are cell cycle-dependent. Both in G1-arrested and G2/M-arrested cells, end-tethering loss is rare and is unaffected by deleting Mrc1or Ctf4 (Fig. 6C). *mec1*Δ cells also show no defect in end tethering either in cycling cells or after G2/M-arrest. This result implies that Mrc1 and Ctf4-dependent end-tethering may be more important in the absence of Mec1-dependent cell cycle arrest.

### Mrc1 and Ctf4 limit DNA end-resection

Using the same fluorescence assay we also investigated whether Mrc1 or Ctf4 regulate DNA 5’ to 3’ end resection, where loss of one or both fluorescent foci was measured 2.5 h after DSB induction. Consistent with previous results, Mrc1 plays a role in limiting resection at a single DSB (21,20). Loss of one or both foci is significantly elevated in *mrc1*Δ cells compared to wild type; but here, too, Tof1 plays only a minor role. Both *mrc1*Δ869–1096 and *mrc1*Δ355–670 show defects in resection limitation (Fig. 7A). Again, *ctf4*D resembled *mrc1*D and the double mutant showed epistasis.

**Figure 7.**
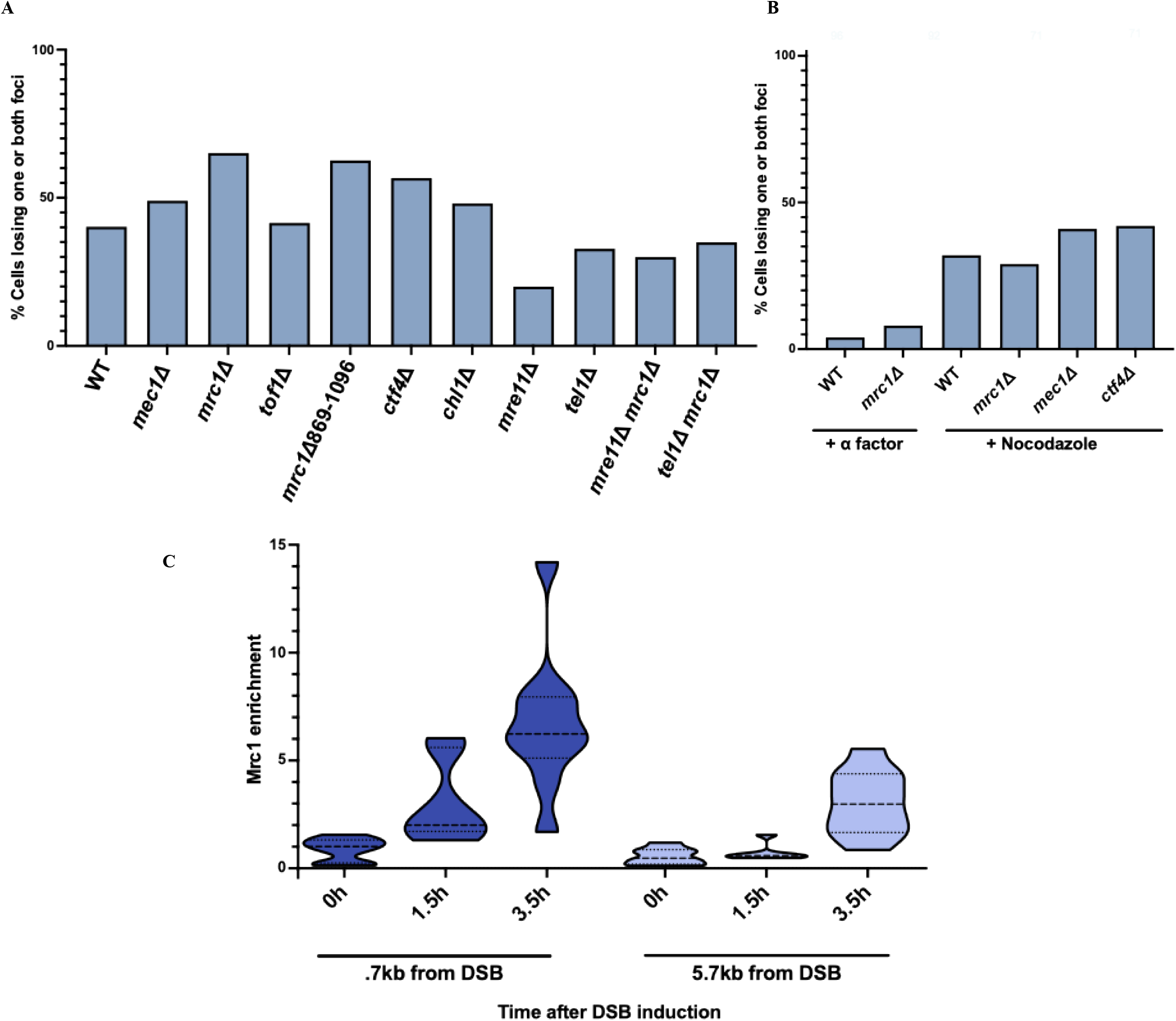
Mrc1 limits end resection during S-phase dependent on canonical resection factors and accumulates near the DSB. A. Quantification of resection, measured by loss of one or both fluorescent foci. The increased resection in *mrc1*Δ cells is suppressed by deletion of Fun30, Mre11, or Tel1. B. *mrc1*Δ resection limitation defect is abolished by G1 arrest (alpha factor) or G2/M arrest (nocodazole). C. ChIP of Mrc1 at 700 bp and 5 kb from the DSB, y-axis is fold Mrc1 enrichment normalized to an uncut locus and to input sample. Median shown in dashed horizontal line, quartile shown in dotted horizontal line.

The *mrc1*Δ resection phenotype is suppressed by slowing the rate of resection through deletion of either Mre11 or the chromatin remodeler Fun30 (Fig. 7A). Tel1 similarly contributes to resection at the DSB: *mrc1*Δ *tel1*Δ double mutants show reduced resection relative to *mrc1*Δ single mutants, suggesting that Tel1 regulates resection, likely through the MRX complex (7). As with end tethering, Mrc1 and Ctf4’s resection-limiting function is observed primarily during S phase, with minimal effects during G1 or G2/M arrest (Fig. 7B).

To determine whether Mrc1 promotes DSB end tethering and resection limitation through direct interaction at the break, we investigated its localization by ChIP-qPCR. Mrc1 accumulates at DSB sites with enrichment at both 0.7 kb and 5 kb from the break, demonstrating direct association with DSB-proximal chromatin (Fig. 7C).

Replication origins (ARS) are distributed across all chromosomes and fire with defined timing during S phase, such that a DSB occurring early in S phase will be engaged by forks fired from the nearest origins. Deleting one of these proximal origins is therefore expected to reduce the number of stalled forks that accumulate at the break site, particularly in the early S phase when distal origins have not yet fired. We found a reduction in Mec1-independent survival when a DSB-proximal, early firing ARS, ARS306, is deleted in strain tNS2333 (Fig. S7). While chromosome III harbors many origins where replisomes are loaded and replication is initiated, loss of one near the DSB site might result in reduced and delayed accumulation of replisome-associated factors at the DSB. Overall, this implies that Mrc1’s role may depend on local accumulation of replication forks near the break during S phase.

### Mrc1, but not Tof1 or Csm3 is required for normal heterochromatic silencing

Mrc1 has been implicated in heterochromatic gene silencing in both fission yeast and budding yeast (39,40,34). To characterize this regulation further, we examined silencing at the *HML* locus in budding yeast using the CRASH assay (41). This assay measures transient unsilencing of the CRE recombinase gene inserted into the silent mating-type locus, *HML*. If *HML::CRE* is even transiently unsilenced, CRE will be expressed and cause a deletion within a reporter construct, irreversibly switching the cell and its descendants from expressing RFP to GFP. We confirmed that Mrc1 is required for normal heterochromatic silencing of *HML* as well as *HMR* (34), with the great majority of *mrc1*Δ colonies switching from red to green; however, this silencing function is independent of Mrc1’s partners, Tof1 and Csm3, which have only a minor effect (Fig. 8). Similarly, we recapitulate findings from Yu et al. (2024), reporting that *mrc1*Δ cells lose silencing at telomeres (Fig. S8); here, Tof1 has a mild impact on telomeric silencing. This result parallels Mrc1’s independence of Tof1 and Csm3 in DSB repair, emphasizing that Mrc1 performs several functions that are distinct from its canonical replication fork checkpoint partners.

**Figure 8.**
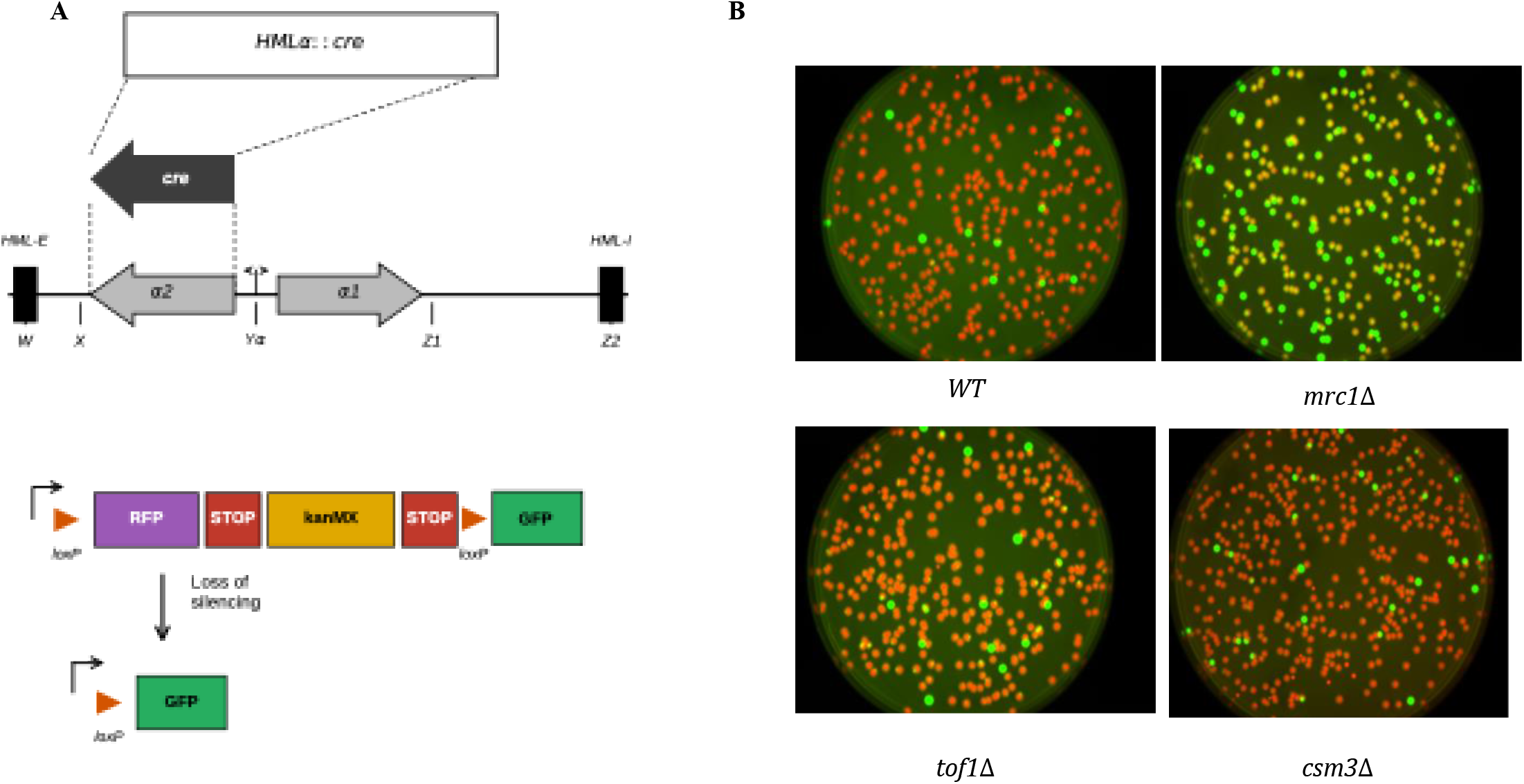
**Mrc1 promotes heterochromatic silencing at HML independently of Tof1 and Csm3**. A. Schematic of CRASH system to measure silencing (adapted from Dodson and Rine, 2015). B. Colony images of wild-type, *mrc1*Δ, *tof1*Δ, and *csm3*Δ cells. Red sectors indicate cells maintaining HML silencing; green indicate cells that have permanently lost silencing. *mrc1*Δ colonies show markedly more green sectors than wild type, *tof1*Δ, or *csm3*Δ, indicating more frequent loss of HML silencing

## Discussion

Our findings establish a checkpoint arrest-independent pathway that enables DSB survival in the absence of Mec1 and reveal unexpected roles for Mrc1 in coordinating DSB repair.

The Tel1/Rad53/9-1-1 pathway we describe enhances repair in the absence of checkpoint-mediated arrest that normally allows cells sufficient time to repair a DSB without entering mitosis. We show that when Mec1 is absent, the 9-1-1 complex independently supports Tel1 accumulation at DSBs, and Tel1 supports DSB end-tethering with Mre11, enhancing repair and survival. It must be noted that, despite both kinases phosphorylating S/TQ sites in many of the same proteins, Tel1 does not simply substitute for Mec1, as deletion of Mec1 completely abolishes G2/M arrest in response to a single DSB. However, Tel1 has been shown to phosphorylate sites on Rad53 in the absence of Mec1 (42), which may account for Tel1’s residual role in Mec1-independent repair.

The finding that Rad53’s role in this context is kinase-independent suggests that Rad53 has structural or alternative functions normally masked by canonical Mec1-dependent phosphorylation. Rad53 also facilitates binding to histones H3-H4 and promotes their degradation to maintain proper chromatin configuration (43) — a function that could facilitate nucleosome reassembly during the extensive new DNA synthesis required for BIR,however, this histone management role requires Rad53 kinase activity, and since kinase-dead Rad53 only mildly impacts Mec1-independent survival, it is more likely thestructural chromatin association of Rad53, rather than active phosphorylation that is relevant in this context. Further experiments will need to be done to understand how Rad53’s kinase activity contributes to DSB survival in this context. Another striking feature of this Rad9-independent, Rad53 kinase-independent pathway is that Mrc1 activity is independent of its S/TQ phosphorylation sites. However, Tel1 independently phosphorylates Mrc1 at several other sites (44,45). Future work will determine whether Tel1 phosphorylation of these sites plays a critical role.

A notable consequence of checkpoint arrest-independence in this pathway is that *mec1*Δ cells progress into mitosis carrying an unrepaired or partially repaired DSB. In this context, end tethering takes on particular significance beyond its established role in facilitating repair and homology search: maintaining physical proximity between broken chromosome ends could preserve the structural continuity of the broken chromosome, promoting efficient repair in cells without installed arrest. The correlation between tethering proficiency and Mec1-independent survival we observe is consistent with the idea that tethering could provide a physical connection. Both pedigree analysis of the pattern of repair in *mec1*D cells (9) and more recent monitoring of a fluorescently-tagged acentric fragment (46) argue that the sister chromatid copies of an acentric fragment remain tethered to each other and do not segregate at mitosis.

An important caveat in interpreting repair pathway choice in this system concerns the relationship between DSB induction and cell cycle position. Because HO endonuclease is induced in asynchronous cultures, individual cells experience the break at different cell cycle stages. However, given that roughly 80% of cycling yeast cells are in G1 or S phase at any given time, the majority of cells in our experiments are likely experiencing the break prior to or during replication. If cleavage occurs during replication, a replication fork may encounter the DSB, generating a one-ended break that is a preferred substrate for BIR rather than SSA. If cleavage occurs after replication, both sister chromatids may be cleaved simultaneously, eliminating the intact sister as a repair template. Cell cycle heterogeneity likely contributes to the variable survival and repair kinetics we observe across mutants.

The interplay between BIR and SSA is difficult to dissect. For example, depletion of Rad51 may hinder BIR while encouraging SSA, and vice versa for *fun30*Δ. Although *mrc1*Δ leads to increased resection, which may benefit SSA and inhibit BIR, a *mec1*Δ *mrc1*Δ *fun30*Δ double mutant or *mec1*Δ *mrc1*Δ *fun30*Δ triple mutant shows minimal impact.

In strain tNS2333 that we have employed in many of the experiments reported here, both SSA and BIR yield the same outcome. However, it should be noted that in BIR, the formation of a deletion occurs in a way that – at least for a while – leaves the ∼90 kb acentric portion of the broken chromosome – used as a template for BIR – still present (Fig. S1). The presence of this long, broken fragment may influence recovery after repairing the DSB.

Mrc1 has been characterized mostly as a component of the replication fork checkpoint complex with Tof1 and Csm3. Here we show that Mrc1 has several specific functions distinct from those of Tof1 and Csm3, including promotion of DNA end-tethering at DSBs, limitation of DSB resection, repair by BIR, and heterochromatic gene silencing at HML, HMR, and telomeres. The separation of function between Mrc1 and Tof1 is consistent with prior work showing that Mrc1, but not Tof1, is required for the normal rate of DNA replication fork progression during S phase (15).

Our data suggests that Mrc1’s C-terminal domain serves as a general platform for chromatin-associated activities beyond replication fork checkpoint signaling. The C-terminal domain of Mrc1 emerges as the critical region for end tethering, resection limitation, and DSB survival. The C-terminus harbors interaction sites for Pol δ and Ctf4, and is predicted to be in spatial proximity to Cdc45, while the central domain (residues 433–457) mediates direct binding to Cdc45 and is required for maintaining Cdc45 at stalled replication forks (17). Repair by BIR and GC depends on Pol δ but does not require Cdc45 (47). It is likely that Mrc1’s interaction with Ctf4 through the C-terminus is important for Ctf4’s role in recruiting Chl1 to the fork, facilitating cohesin binding, and promoting both end tethering and sister chromatid cohesion.

In previous studies, the double deletion of Mrc1 and Ctf4 was reported to be lethal (48). This synthetic lethality of *mrc1*Δ *ctf4*Δ implies that the two proteins have separate but overlapping functions. However, we were also able to recover *mrc1*Δ *ctf4*Δ double mutants in our strain background, albeit slower growing, suggesting this lethality may be strain-dependent. The observation that Mrc1 promotes end-tethering, limits resection, and supports DSB survival is notable in light of parallel findings for Ctf4 and Chl1, which show similar phenotypes at a single DSB. Despite their distinct molecular roles, Mrc1, Ctf4, and Chl1 share a common function in stabilizing cohesin at the replication fork during sister chromatid cohesion establishment. That all three influence DSB end tethering suggests this replisome-cohesin-stabilizing activity may be repurposed at DNA ends — maintaining cohesion or structural organization proximal to the break to support end-tethering.

The domain spanning residues 355–670, which contains predicted cohesin and Cdc45 binding motifs, also shows minor contributions to both end tethering and resection limitation. It is unclear whether Mrc1 directly or indirectly interacts with resection machinery to limit resection progression, but we show that the increased resection seen when Mrc1 is deleted depends on canonical resection pathways that involve Fun30 and Mre11. Loss of Fun30 compromises long-range resection (Exo1 and Sgs1-Dna2), while Mre11 is involved only in initial short-range resection, but still impacts repair kinetics (49). How each affects Mrc1-dependent resection limitation is not yet clear. Importantly, Mrc1’s role in end tethering is separable from its resection limitation function, as seen in *mrc1*Δ *fun30*Δ mutants. Although long-range resection is fully active in checkpoint-arrested G2/M cells (50,51), the effect of Mrc1 appears constrained to S phase.

Future studies to dissect specific molecular interactions within the Mrc1 C-terminus will help elucidate the mechanisms by which it coordinates these functions. It seems likely that both Mrc1’s role in resection limitation and in DSB end tethering are important for preventing lethal genomic rearrangements and for timely and accurate repair, which is especially critical in the absence of Mec1-mediated arrest.

Mrc1’s particular importance for HR is notable, given requirements for homology search, strand invasion, and templated DNA synthesis — processes that may be facilitated by maintaining end-tethering and limiting extensive resection. The Tel1– and Mrc1-dependent, but Mec1-independent, GC efficiency suggests that checkpoint and repair fidelity mechanisms are tightly coordinated even when Mec1 is absent, implying an arrest-independent pathway relying on checkpoint-associated factors to promote DSB repair.

Taken together, these convergent results point to a broader and underappreciated contribution of replication fork-associated machinery to the resolution of a single DSB. The replication stress response is rarely invoked in discussions of DSB repair, where the DNA damage checkpoint rightly dominates.

However, DSBs that arise during S-phase exist in a cellular environment where replication forks are actively engaged across the genome. It is likely that forks encountering a DSB engage the same stabilization machinery that responds to replication stress — and that this fork-associated response promotes end-tethering, limits resection, and may promote repair. Under normal conditions, this contribution is overshadowed by the robust Mec1-dependent DDC arrest. The loss of Mec1, which reveals this pathway, suggests that replication forks are not merely bystanders to DSB repair in S-phase, but active participants in its resolution.

The S-phase specificity of both Mrc1’s end tethering and resection limitation suggests that these functions integrate with the replication machinery or with S-phase-specific repair factors. Whether Mrc1 accumulates at DSBs during S phase due to proximity to normal replication forks or through damage-specific signaling remains to be determined. However, we have previously shown that *ctf4*Δ impairs BIR in nocodazole arrested cells (52). In a broader context, these findings highlight the concept of checkpoint substitution—cells can activate or utilize alternative pathways when canonical checkpoints are unavailable. This idea has important implications for cancer biology, where checkpoint defects are common. Our work suggests that even cells with severe ATR deficiency might retain substantial repair capacity through backup pathways, which could allow survival of otherwise lethal DNA damage.

## Methods

### Yeast strains and plasmids

We often employed strain tNS2333 (8), a derivative of strain YMV80 (10). Gene deletions and epitope tags were introduced by standard one-step PCR-based methods using selectable marker cassettes as previously described (53,54). Centromeric plasmids used for complementation of *rad53*Δ and *mrc1*Δ alleles were introduced to express genes under their native promoters. A complete list of strains and plasmids are provided in the Supplementary Information, Tables 1 and 2. H2AS129 point mutants were made using Cas9 targeting *H2A1* and *H2A2*, and an 80mer introducing the mutations. Oligonucleotides listed used to construct guide RNAs are listed in Supplemental Table 3 with ends complimentary to a BplI site were annealed and ligated into plasmids bRA89 and bRA90 with constitutively expressed Cas9. *Rad53-kd* mutants were also created using a Cas9 and an 80mer introducing the K227A mutation.

### DSB induction and viability assays

DSBs were induced using an integrated galactose-inducible HO endonuclease (55,56). Cells were grown in YP-lactose to mid-log phase and DSB induction was initiated by addition of 2% galactose. For viability assays, cells were plated onto YP-dextrose or YP-galactose and colonies counted after 2 or 3 days at 30°C respectively. Survival was calculated as the ratio of colonies on DSB-inducing conditions (galactose) relative to non-inducing (dextrose) conditions.

### Cell cycle analysis

Cell cycle morphology was assessed by DAPI staining (VECTASHIELD® Antifade Mounting Medium with DAPI; Vector Laboratories, cat. no. H-1200) following fixation in 70% ethanol. Large-budded cells with a single DAPI signal in the bud-neck region were scored as G2/M-arrested. For synchronization, cells were arrested in G1 with 1 µM alpha factor (US Biological, cat. no. N3000) or in G2/M with 20 µg/mL nocodazole (CAS: 31430-18-9) for 3 h before DSB induction.

### PCR assays

PCR assays to quantify repair kinetics were performed as previously described (26). Briefly, strains were grown overnight in YEP-Lac and DSBs were induced by addition of galactose to 2% final concentration. Samples were collected at 0, 3, 6, 9, and 12 hours post-induction, with an additional wild-type sample collected at ∼24 hours as a fully repaired control. Following DNA purification, samples were resuspended in 0.5x TE, quantified by NanoDrop, normalized to 500 ng/µl, and diluted to a working concentration of 40 ng/µl, with further adjustments made based on prior PCR runs. PCRs were performed using 2×EsTaq MasterMix (CWBio, cat. no. CW0690L) with a 2-minute elongation step. Repair products were amplified using primers MA193 and LEU2RP for 28 cycles; control amplicons were generated using primers EB29 and EB30 for 25 cycles. Products were resolved by electrophoresis on 1% agarose gels stained with EtBr and quantified using Image Lab software (Bio-Rad). Quantification was performed against a standard curve of serial 1:2 dilutions of purified 100% repaired DNA, spanning a range of five dilutions calibrated separately for control and repair amplicons. Percentage repair was calculated by dividing the repair signal by the signal from the independent control locus GLC7, then normalizing to the ratio from the 24-hour wild-type sample. At least three independent time-courses were performed per strain, with a minimum of three technical PCR replicates per sample.

### Chromatin immunoprecipitation

ChIP was performed as previously described (57). Briefly, cells were cross-linked with 1% formaldehyde (Sigma-Aldrich, cat. no. 47608) for 15 min at room temperature, quenched with glycine, and lysed by bead beating. Chromatin was sheared by sonication to 200–500 bp fragments. Immunoprecipitation was performed using anti-HA (HA-tag(C29F4) rabbit monoclonal; Cell Signaling Technology, cat. no. 3724) or anti-Myc (mouse monoclonal; Abcam, cat. no. ab16918; RRID: AB_30256) antibodies conjugated to Protein G Dynabeads (Invitrogen, cat. no. 10003D). DNA was quantified by qPCR using primers flanking the DSB, normalized to an uncut locus (*ADH1*) on chromosome XV.

### Microscopy and end-tethering and resection assay

End tethering was visualized using a dual-fluorescence system in which TetR-GFP marks one DSB end and LacI-mCherry marks the other, as described by 28. Cells were imaged on a Nikon Ni-E upright microscope equipped with a Yokogawa CSU-W1 spinning-disk head, an Andor iXon 897U EMCCD camera, Nikon Elements AR software, a 100x oil immersion objective, and a 488 nm and 561 nm laser. Approximately 18 z-stacks with a thickness of 0.2 µm were collected per image. Images were acquired at 2.5 hours post-galactose addition. Images were processed and analyzed using Fiji (NIH). End tethering was scored as the fraction of cells with separate RFP and GFP foci to those with overlapping foci. At least 100 cells were scored per genotype across three independent experiments.

### Silencing assays

CRASH strains were modified from strain JRY968 (41). Strains were grown and imaged as previously reported (58). CRASH strains were pre-selected on G418 to assure retention of the *loxP:RFP:kanMX:loxP:GFP* cassette. Cells were picked up from G418 plates resuspended in water, and serially diluted to reach a density of approximately 100-200 cells per plate, on YEP-dex plates. Colonies were incubated for 4-5 days at 30°C and were then imaged as whole plates using a BioRad Chemidoc MP Imaging System.

## Statistical analysis

Statistical analyses and graphing were performed using GraphPad Prism 7.00 (GraphPad Software, Inc.). All raw data and statistical outputs are provided in Supplementary Tables. CFU plating assays were performed in at least three independent biological replicates; survival ratios were compared between genotypes using a two-tailed Welch’s *t*-test. For microscopy-based end-tethering and resection assays, at least 100 cells were scored per genotype per experiment across a minimum of three independent experiments, and genotypes were compared using Fisher’s exact test. ChIP-qPCR values represent the mean of at least three independent biological replicates (nine technical replicates total), normalized to an uncut control locus. All error bars represent SEM. A *p*-value of < 0.05 was considered statistically significant. Significance is indicated on figures as follows: * *p* < 0.05, ** *p* < 0.01, *** *p* < 0.001, **** *p* < 0.0001. Bonferroni correction was applied for multiple pairwise comparisons in resection data (105 total comparisons); Benjamini-Hochberg (BH) false discovery rate correction was applied for end-tethering comparisons (91 total comparisons). For repair kinetic assays, survival was quantified as area under the curve (AUC) by trapezoidal integration over 0–12 hr; genotypes were first compared using a Kruskal-Wallis test, followed by pairwise exact Mann-Whitney U tests with Benjamini-Hochberg FDR correction (15 comparisons per experiment).

## Data Availability

Data are provided in several Supplementary Data and Tables. Additional data are available upon request to the communicating author.

## Supplementary figure legends

**Figure S1.**
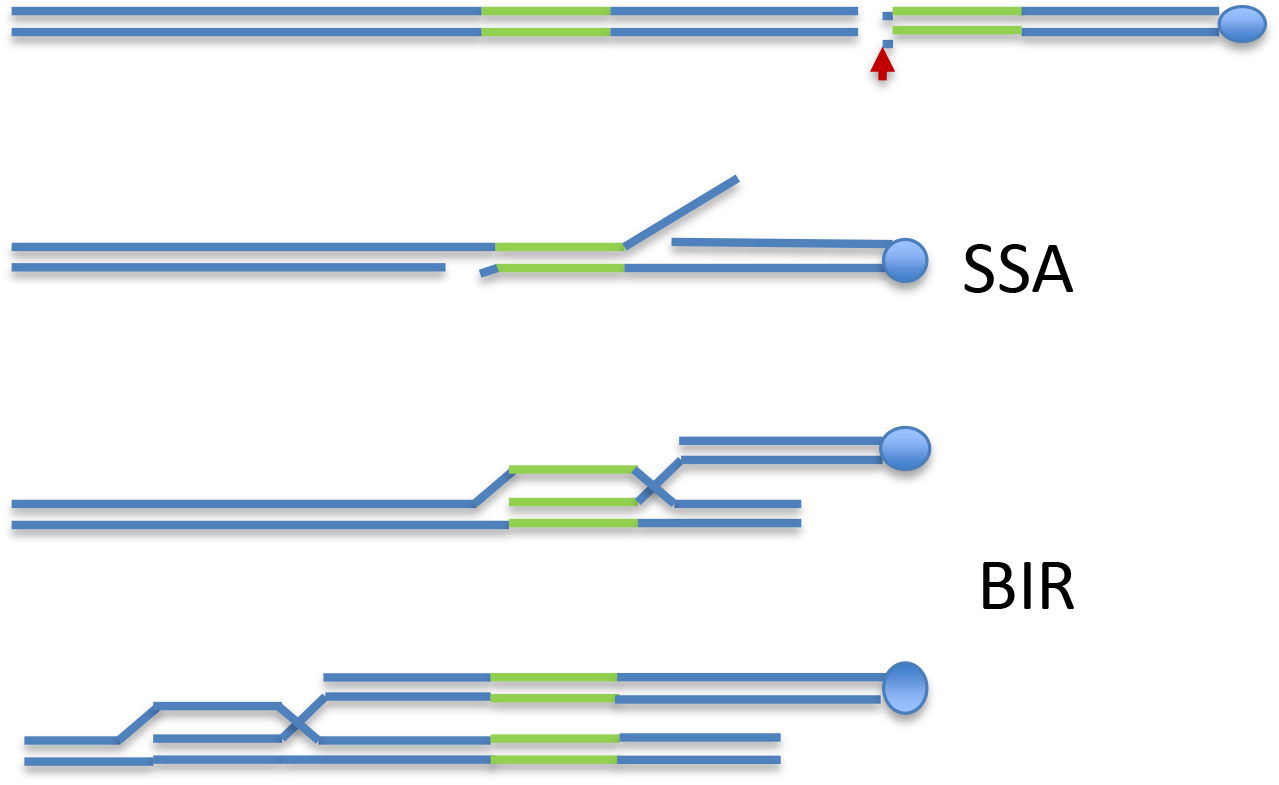
Alternative repair of a DSB in strain tNS2333. tNS233 can be repaired through BIR or SSA. Although both products are the same, BIR will harbor a large, acentric chromosome fragment until it is degraded.

**Figure S2.**
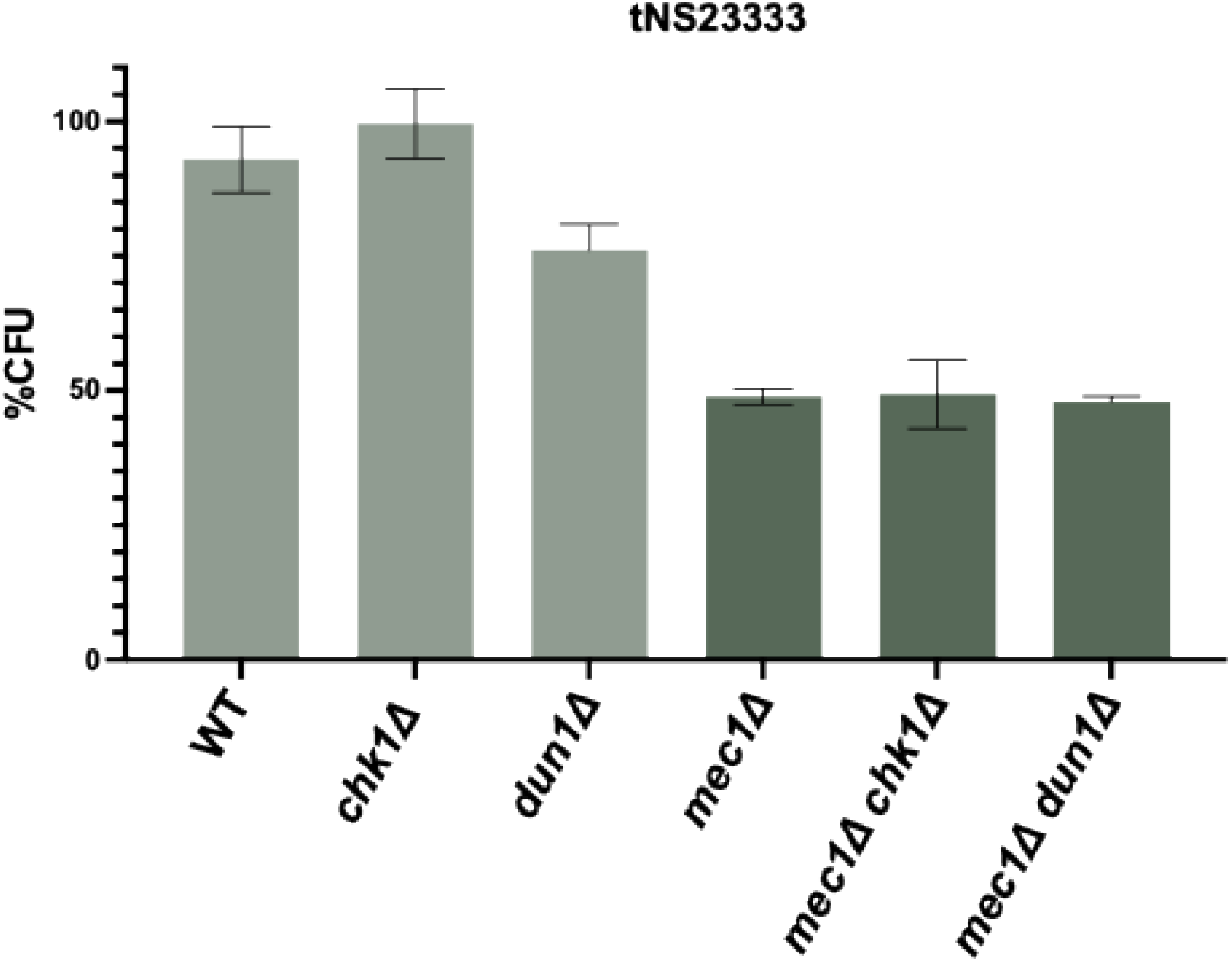
Neither Dun1 nor Chk1 are important for Mec1-independent survival. CFU assay in tNS233, Chk1 and Dun1 have no impact on Mec1-dependent or independent survival.

**Figure S3.**
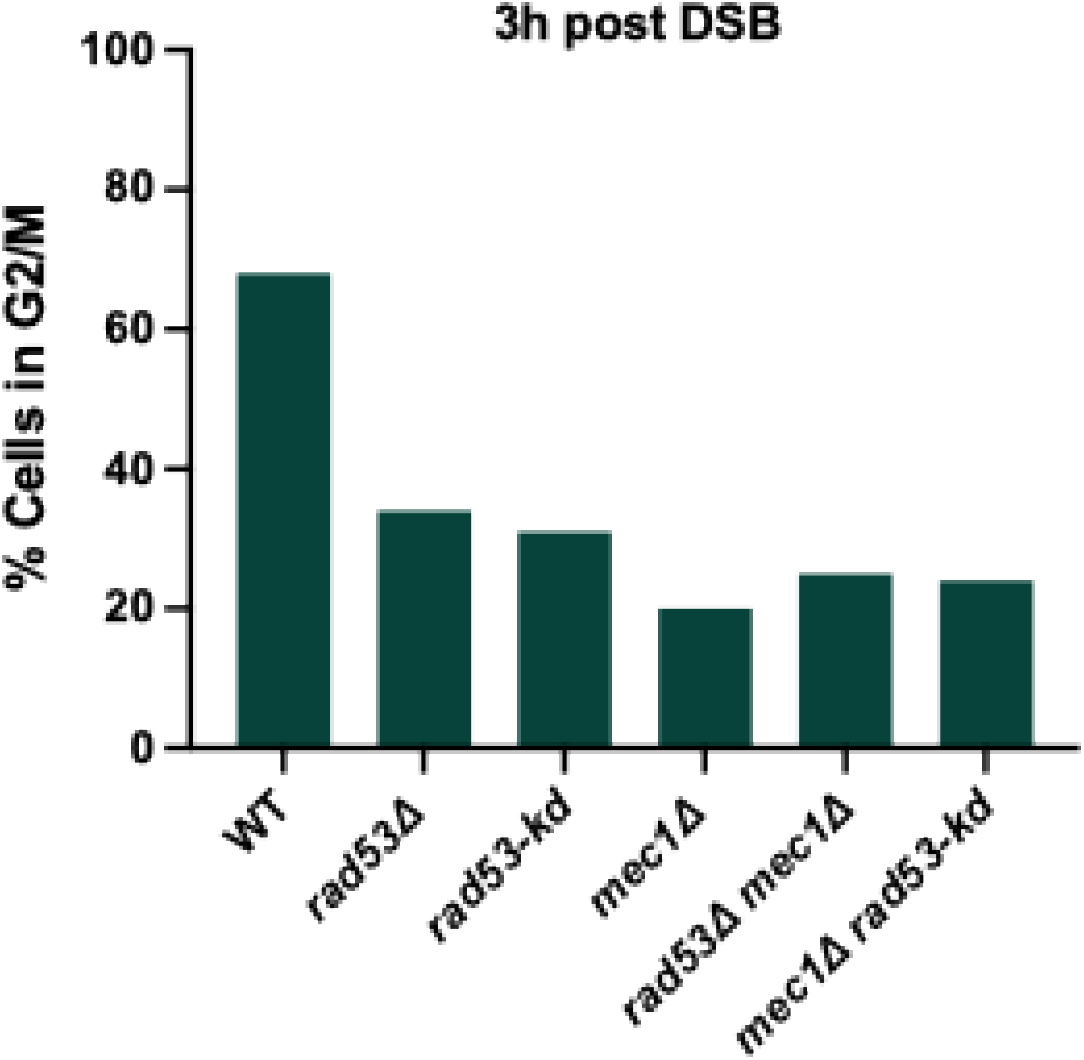
Rad53-kd mutants do not arrest. Percentage of cells in G2/M arrest 3h after DSB induction in tNS2333. >100 cells collected for each mutant.

**Figure S4.**
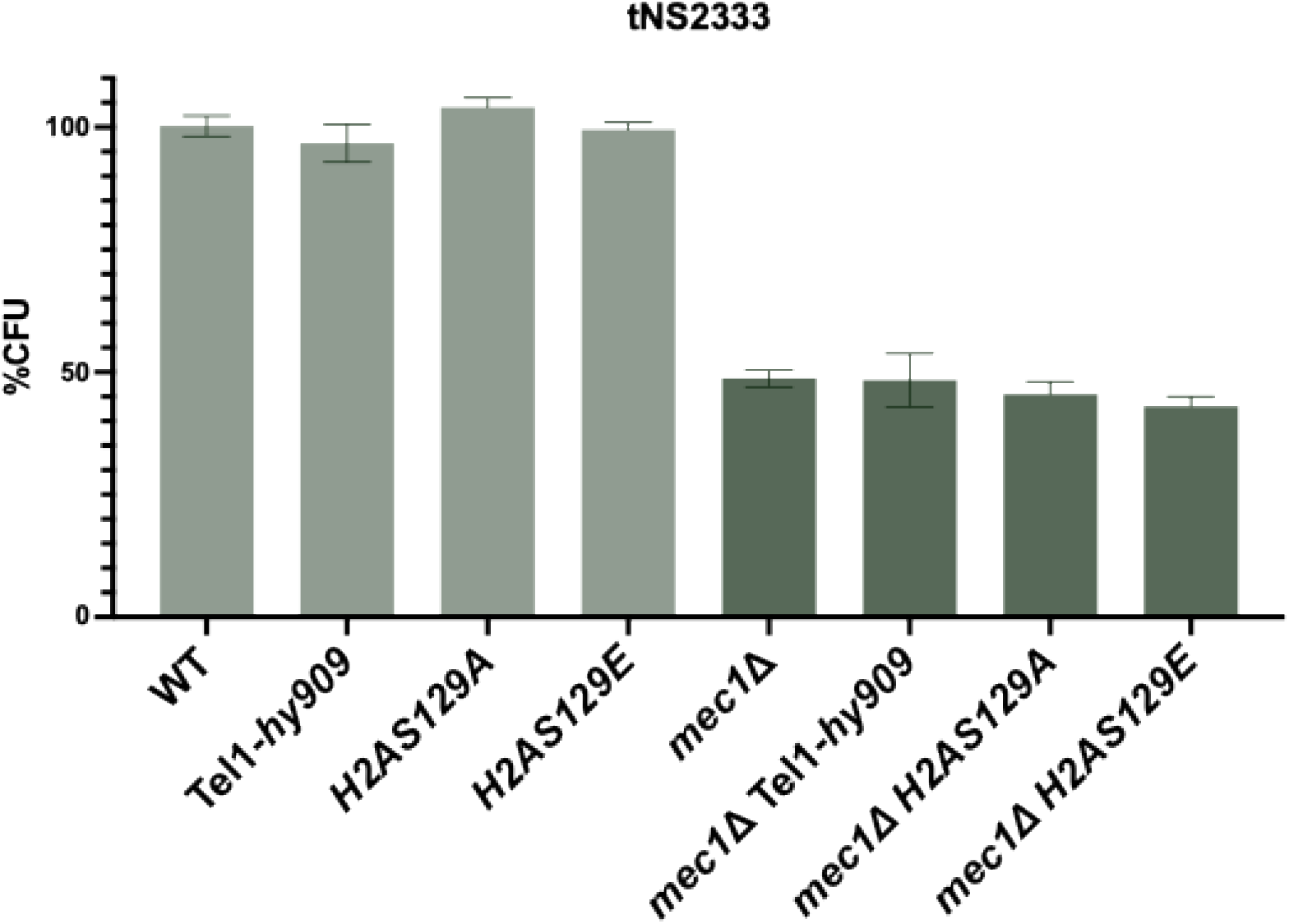
Neither hyperactive Tel1 nor γ-H2A is important for Mec1-independent survival. CFU assay in tNS2333. hyperactive Tel1 mutant tel1-hy909 shows no improvement for Mec1-independent survival. Neither H2A-S129A nor H2A-S129E impact Mec1-independent survival.

**Figure S5.**
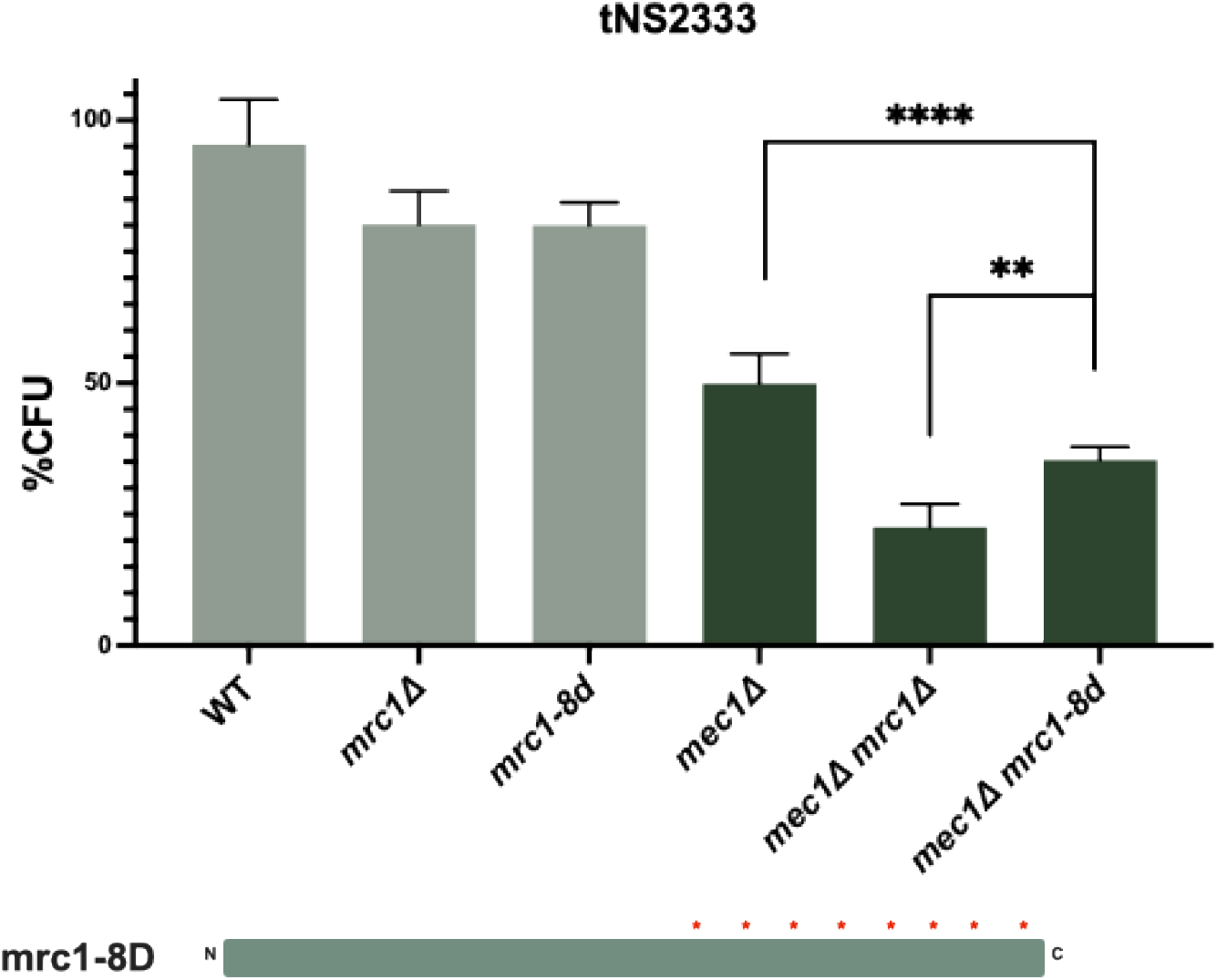
mrc1-8D shows a partial defect. CFU assay in tNS2333, *mec1*Δ and *mrc1-8D* mutants (8 Rad53 phosphorylation S or T sites are converted to D).

**Figure S6.**
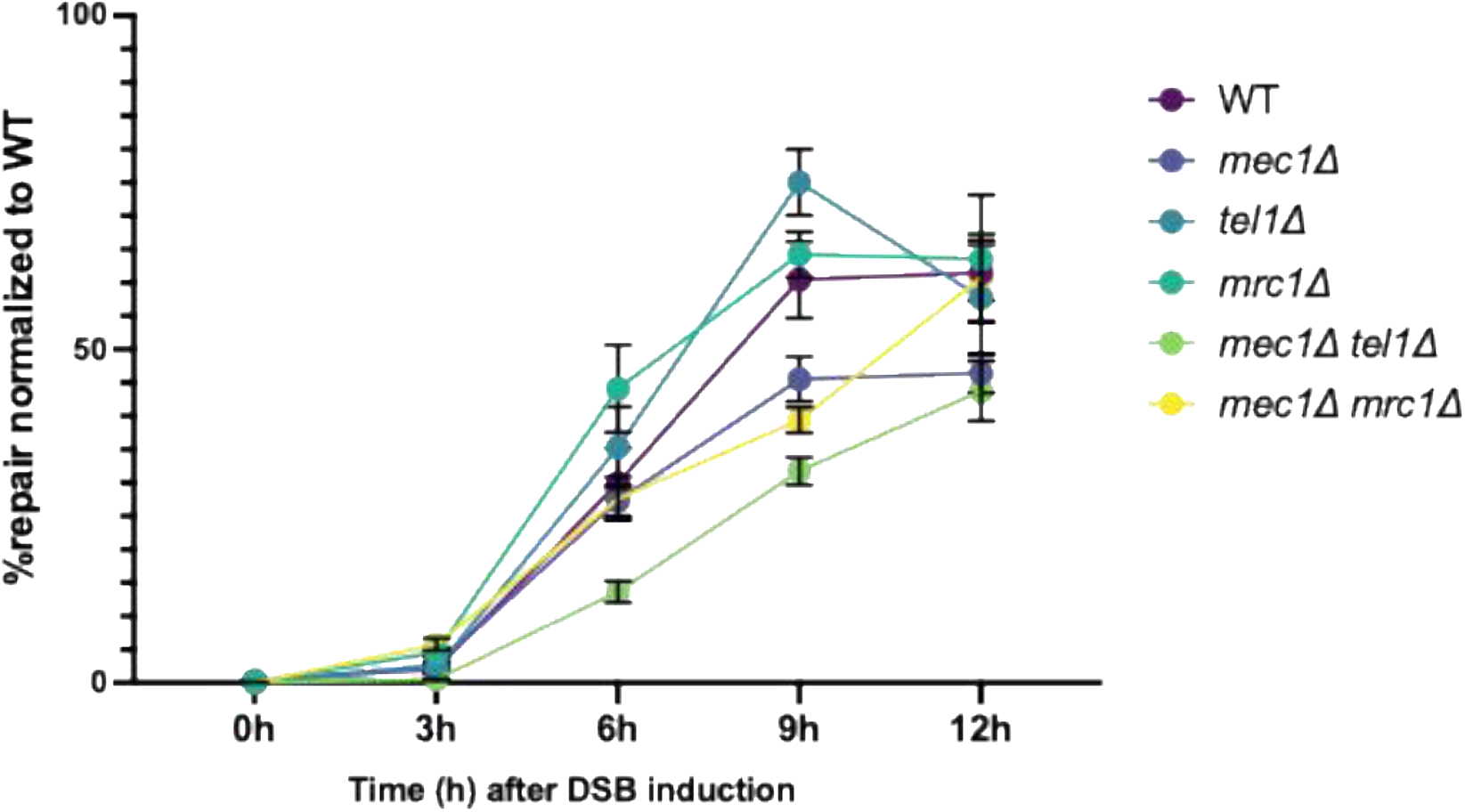
Nocodazole arrested cells show rescue of repair in *mec1*Δ cells. In tNS2333, *mec1*Δ*, mec1*Δ *mrc1*Δ and *mec1*Δ *tel1*Δ all show rescue of repair in cells arrested in nocodazole 3 h before DSB induction. The y-axis is normalized to the percent PCR signal compared to cells that had previously repaired the DSB (WT colony grown on YEP-GAL) and normalized to a control amplicon at *GLC7*.

**Figure S7.**
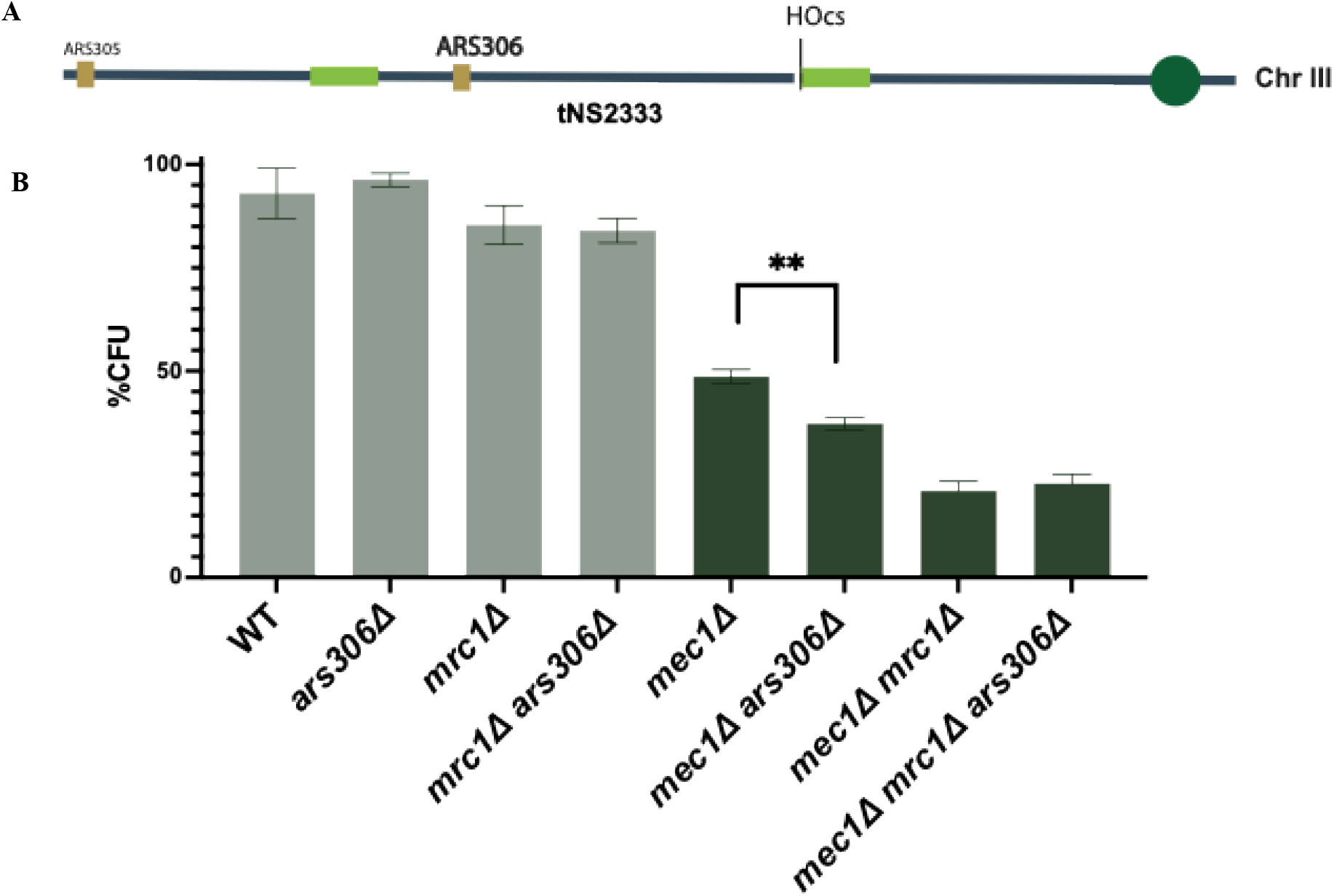
Loss of replication origin ARS306 significantly reduces survival in *mec1Δ* cells. A. diagram of the location of origins of replication on the left arm of chromosome 3. B. CFU assay in tNS2333.

**Figure S8.**
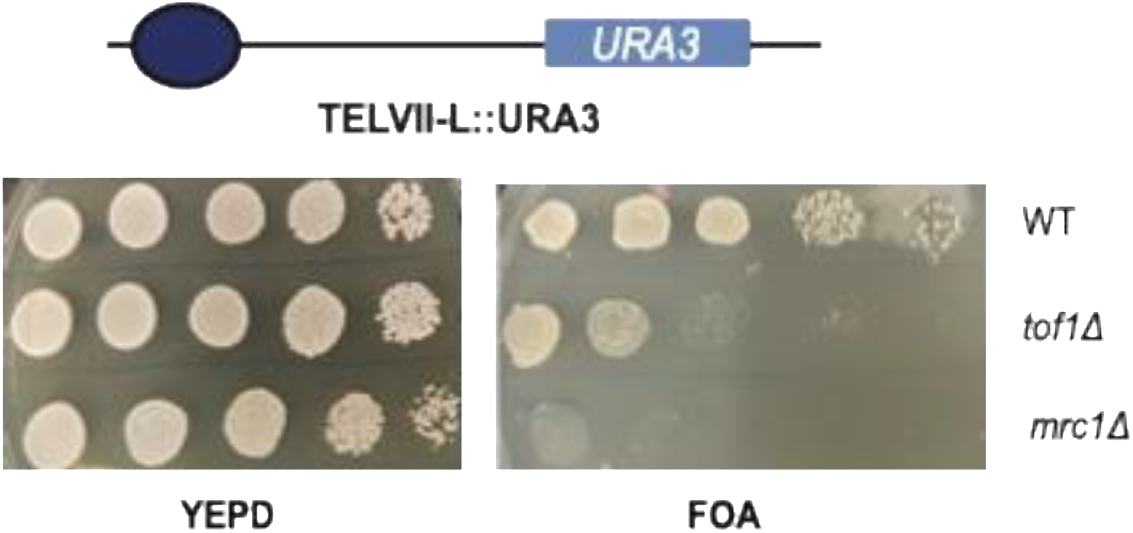
Loss of silencing at telomeres in *mrc1*Δ is more severe than tof1Δ. A. Telomere silencing assay (adapted from Yu et al., 2024) DMY3314, where a URA3 gene is placed at the telomere. B. Cells plated on FOA vs YEPD. Loss of silencing is measure by ability to grow on FOA plates. *mrc1*Δ leads to significantly more loss of silencing at the telomeres than *tof1*Δ.

## Acknowledgments

We thank John Diffley and Steven Elledge for Mrc1 plasmids and Neal Sugawara for his many suggestions and contributions. Manuscript structure and grammatical editing were assisted by Claude (Anthropic).

## Author Contributions

Conceptualization and initial data collection was conducted by M.A. and J.E.H. Data collection, strain making, imaging, primer design, and data analysis were performed by M.A. and E.B. The manuscript was authored and edited by M.A. and J.E.H.

## Data Availability Statement

All data supporting the findings are included in the main text and Supplementary materials. Supplementary Data are available at NAR Online

