## Supplementary data for "Replication Fork-Associated Checkpoint Mediator Mrc1/Claspin Acts in Arrest-Independent Double-Strand Break Survival"

Data for Figure 1d

DSB Repair AUC Analysis — Exact Mann-Whitney U + BH/FDR Correction

AUC per Biological Replicate (trapezoidal integration, 0–12 hr)

| Strain | Genotype | A1 | A2 | A3 | A4 | A5 | A6 | A7 | n | Mean AUC | SD | SEM |
|---|---|---|---|---|---|---|---|---|---|---|---|---|
| MA75 (WT) | WT | 5.90 | | 5.25 | 4.98 | 20.69 | 4.96 | 7.77 | 6.00 | 5.21 | 2.82 | 1.15 |
| MA76 (mec1) | mec1Δ | 2.28 | 1.26 | 1.34 | | 0.72 | 1.23 | 1.06 | 6.00 | 1.89 | 0.29 | 0.12 |
| MA83 (mec1 tel1) | mec1Δ tel1Δ | 1.76 | 1.10 | 1.11 | 0.72 | 0.95 | 1.33 | 0.90 | 7.00 | 0.91 | 0.21 | 0.08 |
| MA84 (tel1) | tel1Δ | 11.72 | | 15.03 | 12.38 | 16.79 | 6.75 | 6.60 | 6.00 | 7.04 | 3.28 | 1.34 |
| MA126 (mc1) | mrc1Δ | 6.16 | | 16.55 | 16.67 | 11.31 | 12.76 | 4.34 | 6.00 | 8.12 | 3.64 | 1.49 |
| MA127 (mrc1 mec1) | mec1Δ mrc1Δ | 1.56 | 1.31 | 1.05 | 1.00 | 1.40 | 1.35 | 0.75 | 7.00 | 1.27 | 0.39 | 0.15 |

Overall Test — Kruskal-Wallis

| | |
|---|---|
| H statistic | 31.9956 |
| Degrees of freedom | 5 |
| p-value | 0.000006 |
| Significant (α=0.05) | **Yes** |

Pairwise Comparisons — Exact Mann-Whitney U + Benjamini-Hochberg FDR correction (15 tests)

| Strain 1 | Strain 2 | Mean 1 | Mean 2 | Δ Mean | U statistic | p (exact) | p (BH adj) | Significance |
|---|---|---|---|---|---|---|---|---|
| WT | mec1Δ | 5.207 | 1.888 | -3.319 | 0.000 | 0.002 | 0.003 | ** |
| WT | mec1Δ tel1Δ | 5.207 | 0.906 | -4.301 | 0.000 | 0.001 | 0.003 | ** |
| WT | tel1Δ | 5.207 | 7.042 | 1.835 | 12.000 | 0.394 | 0.422 | ns |
| WT | mrc1Δ | 5.207 | 8.117 | 2.910 | 9.000 | 0.180 | 0.207 | ns |
| WT | mec1Δ mrc1Δ | 5.207 | 1.267 | -3.940 | 0.000 | 0.001 | 0.003 | ** |
| mec1Δ | mec1Δ tel1Δ | 1.888 | 0.906 | -0.982 | 0.000 | 0.001 | 0.003 | ** |
| mec1Δ | tel1Δ | 1.888 | 7.042 | 5.154 | 0.000 | 0.002 | 0.003 | ** |
| mec1Δ | mrc1Δ | 1.888 | 8.117 | 6.229 | 0.000 | 0.002 | 0.003 | ** |
| mec1Δ | mec1Δ mrc1Δ | 1.888 | 1.267 | -0.622 | 2.000 | 0.005 | 0.006 | ** |
| mec1Δ tel1Δ | tel1Δ | 0.906 | 7.042 | 6.136 | 0.000 | 0.001 | 0.003 | ** |
| mec1Δ tel1Δ | mrc1Δ | 0.906 | 8.117 | 7.211 | 0.000 | 0.001 | 0.003 | ** |
| mec1Δ tel1Δ | mec1Δ mrc1Δ | 0.906 | 1.267 | 0.361 | 8.000 | 0.038 | 0.047 | * |
| tel1Δ | mrc1Δ | 7.042 | 8.117 | 1.075 | 15.000 | 0.699 | 0.699 | ns |
| tel1Δ | mec1Δ mrc1Δ | 7.042 | 1.267 | -5.775 | 0.000 | 0.001 | 0.003 | ** |
| mrc1Δ | mec1Δ mrc1Δ | 8.117 | 1.267 | -6.851 | 0.000 | 0.001 | 0.003 | ** |

### Notes

*AUC = Area Under the Curve via trapezoidal rule (0, 3, 6, 9, 12 hr). Replicates with <3 valid timepoints excluded.*

*Kruskal-Wallis: non-parametric one-way test on AUC ranks. Significant overall effect justifies pairwise tests.*
*Exact Mann-Whitney U: uses the exact permutation distribution (not normal approximation) — critical for small n (6–7 replicates).*

*BH/FDR (Benjamini-Hochberg): controls false discovery rate; less conservative than Bonferroni; appropriate for exploratory multi-strain comparisons.*

*Significance (post-BH correction): *** p<0.001  ** p<0.01  * p<0.05  ns = not significant.*
*ΔMean = Mean AUC of strain2 – strain 1 (positive = strain 2 repaired more, overall.*
