## Supplementary Tables 1-3 for "Replication Fork-Associated Checkpoint Mediator Mrc1/Claspin Acts in Arrest-Independent Double-Strand Break Survival"

Supplemental Table 3

| Oligonucleotide | sequence | purpose | experiment |
|---|---|---|---|
| Ma315mrc1gRNA2fwd | ACTACCAGATGATTCATAATgtttt | to delete 355-670 aa of mrc1 | |
| Ma316mrc1gRNA2rev | ACTACCAGATGATTCATAATgtttt | to delete 355-670 aa of mrc1 | |
| Ma317mrc1gRNA3fwd | ACTACCAGATGATTCATAATgtttt | to delete 711-850 or 798 aa of mrc1 | |
| Ma318mrc1gRNA3rev | ACTACCAGATGATTCATAATgtttt | to delete 711-850 or 798 aa of mrc1 | |
| Ma319mrc1gRNA4fwd | ACTACCAGATGATTCATAATgtttt | to delete 711-850 or 798 aa of mrc1 | |
| Ma320mrc1gRNA4rev | ACTACCAGATGATTCATAATgtttt | to delete 711-850 or 798 aa of mrc1 | |
| Ma321mrc1gRNA5fwd | ACTACCAGATGATTCATAATgtttt | to delete 869-1096 aa of mrc1 | |
| Ma322mrc1gRNA5rev | ACTACCAGATGATTCATAATgtttt | to delete 869-1096 aa of mrc1 | |
| | | | Repair PCR |
| LEU2RP | ACTACCAGATGATTCATAATgtttt | Downstream and antisense to *LEU2*. | Repair PCR |
| MA193_trpfwdforrepair | ACTACCAGATGATTCATAATgtttt | fwd primer at the end of TRP for repair in tns2333 | Repair PCR |
| EB29GLC7ctrlfwd | ACTACCAGATGATTCATAATgtttt | fwd primer for ctrl | Repair PCR |
| EB30GLC7ctrlrev | ACTACCAGATGATTCATAATgtttt | rev primer for ctrl | ChIP experiments |
| MA43ADH1fwd | ACTACCAGATGATTCATAATgtttt | *ADH1* fwd for resection assay ctrl | ChIP experiments |
| MA44ADH1rev | ACTACCAGATGATTCATAATgtttt | *ADH1* rev for resection assay ctrl | ChIP experiments |

| | | | |
|---|---|---|---|
| MA018MATAlphaFWD | ACTACCAGATGATTCATAATgtttt | *MAT*$\alpha$ FWD primer Immediately distal to HOcs | ChIP experiments |
| MA19MATalpharevRSA1 | ACTACCAGATGATTCATAATgtttt | *MAT*$\alpha$ REV primer Immediately distal to HOcs. | ChIP experiments |
| MA36_726bpdownstreamofHOcsfwd | ACTACCAGATGATTCATAATgtttt | 726bp downstream HOcs fwd | ChIP experiments |
| MA37_726bpdownstreamofHOcsrev | ACTACCAGATGATTCATAATgtttt | 726bp downstream HOcs rev | ChIP experiments |
| MA38_5_7kbdownstreamofHOcsfw | ACTACCAGATGATTCATAATgtttt | 5.7 kb downstream HOcs fwd | ChIP experiments |
| MA39_5_7kbdownstreamofHOcsrev | ACTACCAGATGATTCATAATgtttt | 5.7 kb downstream HOcs rev | |
| GM335 | ACTACCAGATGATTCATAATgtttt | 80mer Rad53 K227A | |
| GM336 | ACTACCAGATGATTCATAATgtttt | | |
| GM337 | ACTACCAGATGATTCATAATgtttt | | |
